# Reversible chromatin remodeling enables *Prosopis cineraria* survival under recurrent heat extremes

**DOI:** 10.64898/2026.07.17.734425

**Authors:** Bhumika Dubay, Ramvannish Muthukumar, Fayas Thayale Purayil, Deepak Karthik, Iltaf Shah, Tamilarasan Rajendran, Martin Kottackal, Mohammad Al Oun, Khaled M Hazzouri, Khaled MA Amiri

**Affiliations:** Khalifa Center for Genetic Engineering and Biotechnology, KCGEB, Al Ain, Abu Dhabi, UAE; United Arab Emirates University, Chemistry Department, UAEU, Al Ain, Abu Dhabi, UAE; United Arab Emirates University, Biology Department, UAEU, Al Ain, Abu Dhabi, UAE

## Abstract

Recurrent seasonal heat and drought raise fundamental questions about how long-lived desert plants sustain physiological function across temperature extremes. We have used seasonal profiling at six time points with multilayered omics studies (Hi-C, transcriptomic, histone marks, and DNA methylation) to understand how Prosopis cineraria, a native Arabian desert legume tree, responds to different temperatures and the underlying mechanisms. A clear pattern emerges during peak heat. chromatin boundaries are selectively weakened, and candidate topological domains merge, activating clusters of heat-protective genes that gain active promoter and enhancer marks (H3K4me3 and H3K27ac). In the cool season, immune and developmental gene regulation is coupled with flowering, consistent with a temporal risk-strategy that shifts reproduction away from lethal heat. At the same time, promoter CHH methylation near transposable elements, together with reduced active promoter and enhancer marks (H3K4me3/H3K27ac), points to a proactive developmental phase rather than just surviving the stress. Integrating physiological data, we connect chromatin activation to an SA-ABA reciprocal seasonal profile, MIZ1-associated hydrotropism, and Stay-Green-mediated delayed senescence through chlorophyll retention. With Landscape genomics and phylogenetics, we further identified a housekeeping PEPC with a high predicted melting temperature that could sustain a malate-derived carbon supply, buffering metabolism under heat. Together, these findings reveal that reversible epigenetic gating enables desert trees to survive and recover from extreme seasonal stress.

## Introduction

One of the major challenges for plants in arid regions is surviving recurrent heatwaves, which are becoming more frequent, intense, and prolonged with global heat imbalance. Repeated exposure to extreme heat can suppress photosynthetic CO₂ assimilation, increase respiratory carbon losses, and shift plants toward a negative carbon balance, while also disrupting flowering, pollen performance, fertilization, and seed set, ultimately reducing reproductive success and threatening plant survival[1]. In hyper-arid regions, intense heat often arrives alongside atmospheric drought, leaving only brief periods when conditions are favorable [1]. As a result, perennial plants must pause growth during the harshest months yet be ready to bounce back as soon as the environment allows. Recent crop-model analyses predict that each degree Celsius of warming could cut wheat yields by around 6% [2] unless major adaptations are made [3]. This highlights a fundamental biological dilemma: while resilience mechanisms help protect plant tissues during extreme heat, they can also force a trade-off, shifting resources away from growth, defense, and reproduction.

Most mechanistic knowledge of heat adaptation comes from crops and annual models [1]. Wild perennial relatives preserve strategies shaped by repeated exposure to marginal environments [4, 46, 61]. Desert trees cannot escape heat by completing a life cycle; they must maintain meristems, hydraulics, photosynthetic capacity, and reproductive potential across many summers. Acute heat responses, including heat shock factors and chaperones, contribute to this capacity [5]. However, seasonal survival requires a broader regulatory system able to gate developmental and physiological programs over months and then reverse those states when water and temperature again permit growth.

Chromatin acts as a central regulatory hub for these processes. Active histone marks such as H3K4me3 and H3K27ac help keep genes ready for transcription, while repressive chromatin states can limit developmental programs, and CHH methylation can mark promoters near transposable elements [6,7,14–17, 42]. The three-dimensional organization of the genome adds yet another layer of control: compartments, domains, and loops can shift the accessibility of regulatory regions, especially during stress [8,9,13]. Despite this, very few studies have tracked 3D genome structure, histone marks, DNA methylation, and gene expression together through natural seasonal cycles in a wild perennial.

Here, we use Prosopis cineraria, a native Arabian desert tree legume, to define how seasonal epigenomic plasticity supports survival under repeated thermal extremes. We profiled leaves across February, April, June, August, October, and December, spanning cool, transition, and hot periods, and integrated Hi-C, A/B compartments, insulation boundaries, RNA expression, H3K4me3, H3K27ac, CG/CHG/CHH methylation, phytohormones, pollen viability, root hydrotropism, proteome predicted stability, and landscape genomics from 231 individuals. We hypothesized that P. cineraria uses reversible chromatin remodeling to activate heat-protective programs, repress costly growth and defense modules during peak heat, and restore flowering and recovery programs in cool seasons. The results support this epigenetic-gating model and connect molecular plasticity to water-foraging traits, temporal risk-spreading, candidate metabolic continuity, and adaptive genomic variation.

## Results

### Seasonal heat remodels 3D genome architecture

Seasonal Hi-C maps revealed that hot months did not globally collapse chromatin architecture. Instead, insulation boundaries weakened selectively at reproducible loci, especially boundaries that were strongest in cool seasons, and this weakening coincided with candidate cross-boundary contact gain and candidate topological-domain (TAD)-merge-like behavior across 840 candidate-season comparisons (Fig. 1b-d, Supplementary Fig. 1). Candidate hot-merged domains were enriched for heat/chaperone, ABA-drought-water, ROS/redox, ion-transport, wax/cuticle, immune-receptor and protein-turnover functions, indicating that structural relaxation preferentially affects responsive genomic regions (Fig. 1e). A representative locus on NW_026555344.1 showed hot-season boundary decay near AP2/ERF, WRKY, aquaporin and transporter genes, followed by recovery during October and December (Fig. 1f). Thus, seasonal 3D-genome plasticity appears reversible and constrained rather than damaging (Fig. 1a). Boundary-depth trajectories recovered as the canopy returned to cool-season physiology, supporting a cycle of loosening, candidate contact expansion and restoration rather than a monotonic stress lesion.

**Fig. 1.**
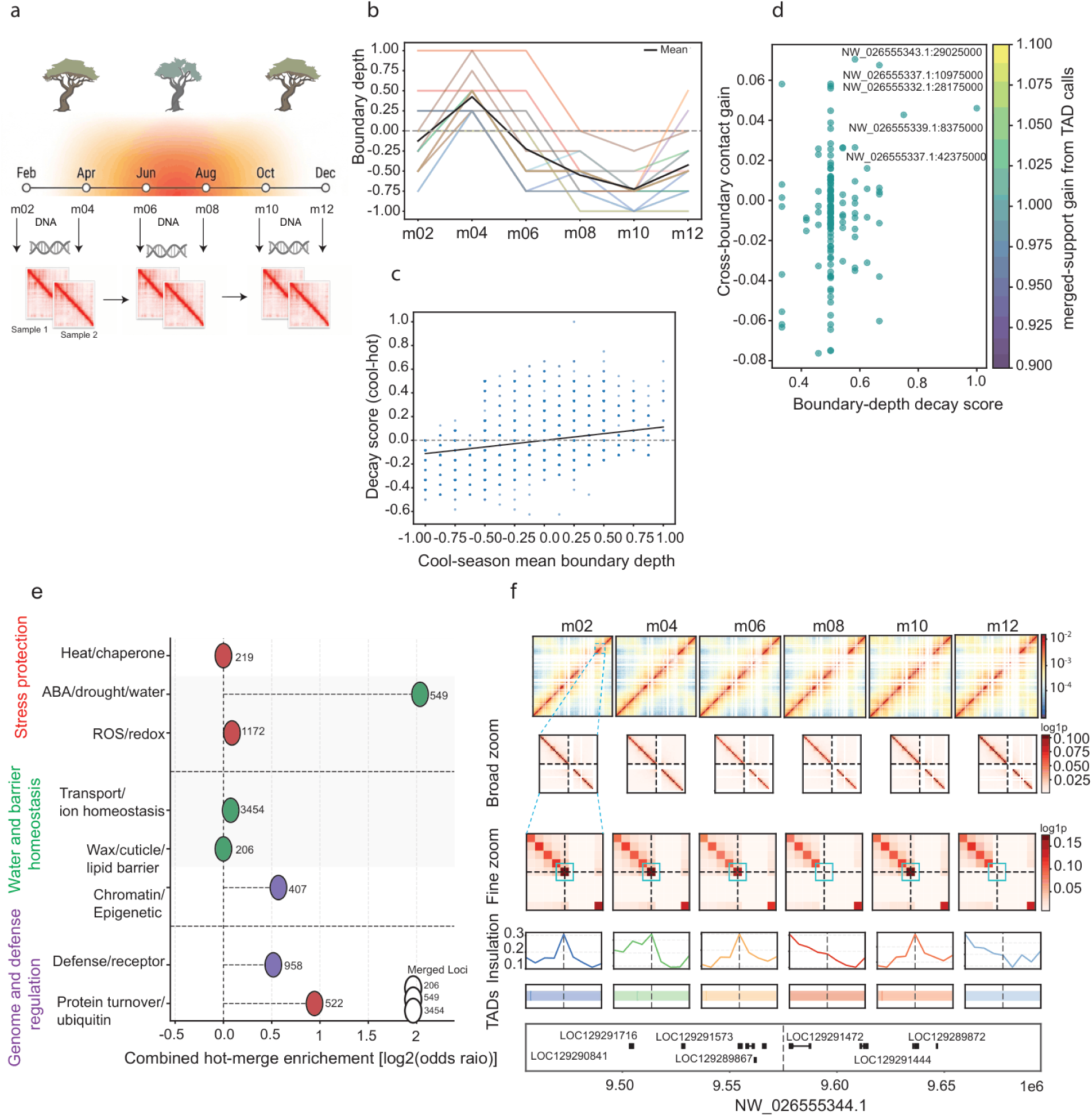
Seasonal heat drives transient 3D genome reconfiguration, boundary-depth decay, and gene-class-specific enrichment within candidate hot-merged domains. **a,** Experimental and analytical design across the seasonal cycle in Prosopis cineraria. Samples collected from cool, transition, and hot months were used to profile chromatin architecture, with representative Hi-C contact maps illustrating seasonal changes in local interaction structure. Hot-season months, particularly June and August, correspond to the strongest heat interval. **b,** Seasonal trajectories of TAD boundary depth across recurrent boundaries. Individual colored lines represent boundary-depth profiles for representative loci, while the black line shows the mean trend across seasons. Boundary depth increases during the early transition into spring but declines through the hot-season interval, consistent with weakening of local domain insulation during peak heat. **c,** Relationship between cool-season mean boundary depth and boundary-depth decay score. The positive regression trend indicates that boundaries with stronger cool-season insulation show greater seasonal decay under hot conditions. **d,** Scatter plot linking boundary-depth decay score with candidate cross-boundary contact gain. Points represent boundary-associated loci, colored by candidate merged-support gain from TAD calls. **e,** Functional enrichment of genes located in candidate hot-merged TADs. Candidate hot-merged domains are enriched for stress-protection categories, including heat/chaperone, ABA/drought/water, and ROS/redox genes, as well as transport/ion homeostasis and wax/cuticle/lipid barrier categories. **f,** Representative locus-level example on scaffold NW_026555344.1 showing seasonal Hi-C contact maps, zoomed contact structure, insulation profiles, and TAD calls across m02-m12.

### Active chromatin coordinates heat-season transcription

Active chromatin marks explained transcriptional seasonality more strongly than accessibility alone. Promoter RNA-H3K4me3 and RNA-H3K27ac correlations were positive across seasons. They strengthened toward the August heat peak (r = 0.54 and 0.50 at m08). In contrast, RNA-ATAC correlations remained weak at all time points, suggesting that accessibility marks regulatory competence rather than productive transcription at individual promoters. CG, CHG and CHH methylation formed a tightly correlated methylation block that was largely independent of RNA, further indicating that methylation contexts co-vary internally but do not systematically predict seasonal expression output (Extended Data Fig. 1). Independent 2025 RNA-seq profiles were concordant with the 2021 multi-omic reference across all seasons (Pearson r = 0.766-0.876; Spearman rho = 0.808-0.904), and expression-bin analysis showed that cross-year agreement extended broadly across the transcriptome rather than being driven only by highly expressed genes (Extended Data Fig. 2). Across 22,587 genes, Cluster 4 (1,331 genes) and Cluster 1 (2,340 genes) were heat-positive, with RNA heat slopes of 12.39 and 7.70 and parallel H3K4me3/H3K27ac gains; the all-gene background showed a much weaker RNA slope of 1.74. Cluster 3 (557 genes) was heat-negative and cool-recovering, yet retained weakly positive active-mark slopes, implying additional repression (Fig. 2a-c, Supplementary Fig. 2, Supplementary Fig. 3). Mann-Whitney and permutation tests supported cluster-specific chromatin gains rather than genome-wide drift. Motif enrichment identified NAC, MYB, WRKY, and ERF/AP2 regulators, and an example of the wax-related nsLTP2 locus showed coordinated hot-season RNA, H3K4me3, ATAC, and H3K27ac activation (Fig. 2d,e). A/B compartment analysis confirmed that active marks and RNA were enriched in the A compartment, especially during the m04-m08 warming transition, with promoter H3K4me3 and H3K27ac A-minus-B differences of 0.07-0.22 and 0.04-0.26 in TSS-centered windows; Spearman correlations between eigenvector-1 scores and assay signals further supported A-like chromatin as a transcriptionally permissive environment (Extended Data Fig. 3, Supplementary Fig. 4).

**Fig. 2.**
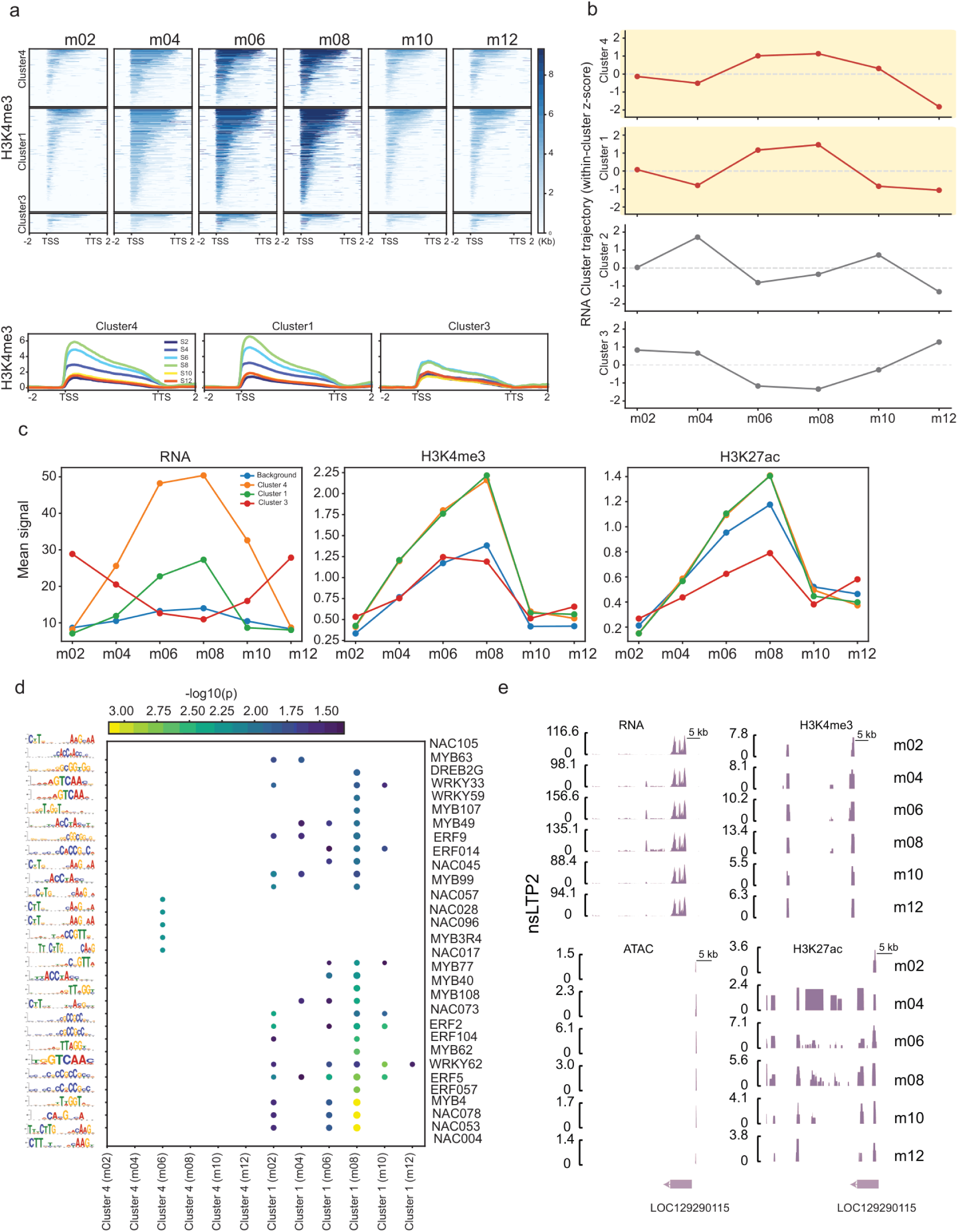
Seasonal coordination of active chromatin, RNA accumulation, and transcription-factor motif architecture across heat and cool-season transitions. **a,** Heat-map and metagene profiles of H3K4me3 signal across six seasonal sampling points. Genes are grouped into four trajectory clusters. Clusters 1 and 4 show progressive enrichment of H3K4me3 toward the hot-season interval. **b,** RNA cluster trajectories plotted as within-cluster z-score-normalized expression across seasons. Heat-positive clusters increase through m06-m08, whereas cool-positive trajectories decline during the hottest months and recover toward m10-m12. **c,** Mean seasonal signal for RNA, H3K4me3, and H3K27ac across clusters and the all-gene background. Coordinated rises in m06-m08 for Clusters 1 and 4 indicate hot-season transcriptional activation accompanied by active promoter/enhancer chromatin marks. **d,** Motif enrichment summary for cluster-associated genes. NAC, MYB, WRKY, ERF, and related stress-responsive regulatory families emerge as candidate regulators of the seasonal chromatin-expression program. **e,** Representative genome-browser tracks at the nsl_TP2 locus (wax-related gene) illustrating seasonal RNA, H3K4me3, ATAC, and H3K27ac co-activation.

### Promoter CHH methylation and low-H3K4me3 states gate seasonal flowering and defense

Methylation and promoter-state analyses identified two conserved promoter-state patterns. CHH methylation was structured across gene bodies and clusters rather than uniformly silencing: heat-positive clusters carried higher CHH signal across much of the year, whereas Cluster 3 showed a hot-season dip followed by recovery (Fig. 3a-c). High CHH with low H3K4me3, especially near the April transition, was consistent with a low-active-mark, transposable-element-associated promoter state; high CHH with high H3K4me3 marked permissive regulatory regions; and low CHH with low H3K4me3 during hot repression suggested reduced chromatin activity at specific photoperiodic targets such as VIN3-LIKE PROTEIN 2 (VIL2) (Extended Data Fig. 4b,e, Supplementary Fig. 5). A broad cool-season low-active-mark set included 1,002 TE-associated promoters dominated by TIR, LTR and Helitron overlaps, spanning ABA/drought, defense, flowering, developmental and chromatin-regulatory genes (Extended Data Fig. 4c). A smaller hot-season low-H3K4me3 module contained 232 promoters during m06/m08, including immune receptors, MADS-box regulators, NF-YB factors and SET/SUVR-like chromatin genes (Extended Data Fig. 4e). Together, these promoter states indicate targeted, context-specific regulation rather than genome-wide promoter silencing, and they require cautious interpretation because CHH methylation can coexist with active regulatory states until direct H3K27me3 ChIP-seq and small-RNA data are available. The cool-up/hot-down program was strongly enriched in Cluster 3 (n = 137; odds ratio approximately 75.7; P = 1.16 × 10^-167) and included disease-resistance and flowering genes that recovered after heat (Fig. 3d,e). GI, CO, SPL, SOC1, FT, and CAL/AP1-like nodes were permissive in February-April and October-December but suppressed in June-August, consistent with cool-season flowering as temporal risk-spreading [26,27] (Fig. 3f, Supplementary Fig. 7).

**Fig. 3.**
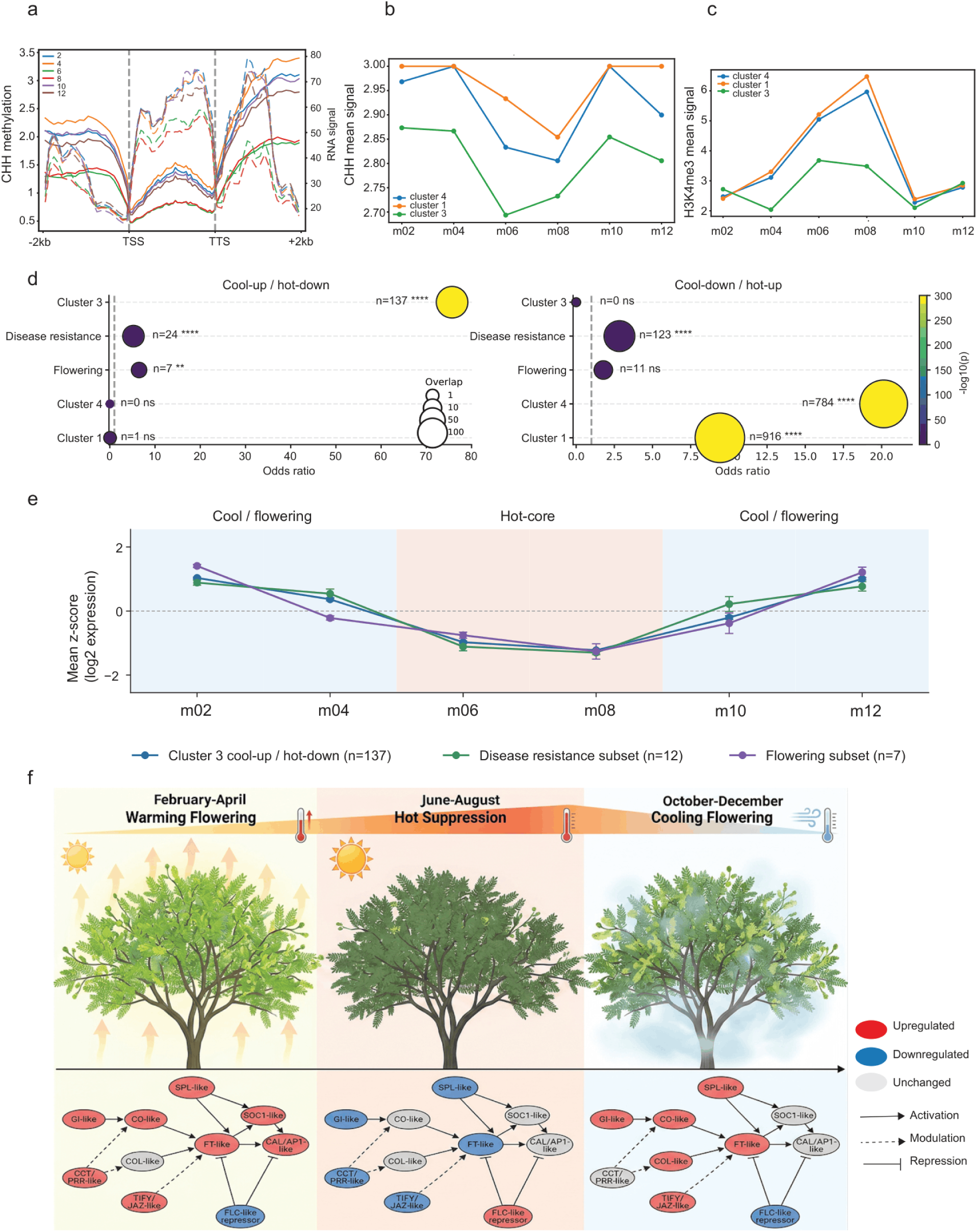
Seasonal CHH methylation and active chromatin define epigenetic gating of cool- and hot-season gene programs. **a,** Metagene profiles showing CHH methylation and RNA signal across annotated gene bodies from 2 kb upstream of the TSS to 2 kb downstream of the TTS across six seasonal collections. **b,** Mean CHH methylation signal across seasonal gene clusters, highlighting cluster-specific methylation trajectories from February to December. **c,** Mean H3K4me3 signal across the same clusters. **d,** Enrichment of cool-up/hot-down and cool-down/hot-up gene sets across seasonal clusters and functional subsets. The cool-up/hot-down program is strongly enriched in Cluster 3 and includes disease-resistance and flowering-associated subsets. **e,** Seasonal expression trajectories of the Cluster 3 cool-up/hot-down module and its disease-resistance and flowering subsets. Mean log2-expression z-scores are high during the cool/flowering windows, decline during the June-August hot core, and recover during October-December. **f,** Working model for seasonal flowering regulation showing permissive states during February-April and October-December and suppression during June-August.

### Compartment switching resolves transition-specific stress pathways

Genome-wide compartment switching further showed transition-specific regulation. Most 50-kb bins remained stable, but reproducible A-to-B and B-to-A switches occurred across adjacent seasonal transitions (Fig. 4a, b). A-to-B switches were enriched for glycosyl hydrolases, ethylene-responsive transcription factors, polygalacturonases, cellulose synthase-like genes, calreticulin, and disease-resistance classes, consistent with attenuation of growth, wall remodeling, and signaling during heat entry (Fig. 4c). B-to-A switches enriched aspartyl proteases, cytochrome P450s, receptor-like kinases, WRKY-related proteins, oligopeptide transporters, expansins, phosphatases, and NADH-dehydrogenase-related genes, indicating activation of detoxification, defense, transport, and remodelling pathways (Fig. 4d). Leaf epicuticular wax peaked in June (15.23 mg g^-1 fresh weight). Across six seasonal means, wax showed a descriptive positive correlation with maximum temperature (Pearson r = 0.63, n = 6) and coincided with candidate distal regulatory interactions involving nsLTP2 and squalene-cyclase-like wax/cuticle genes, supporting a heat-entry programme in which cuticular protection co-occurs with three-dimensional regulatory rewiring (Extended Data Fig. 5). An example cis-regulatory region on NW_026555334.1 contained gamma-glutamyl peptidase-like and oligopeptide transporter genes, linking the same interval to antioxidant and glutathione-associated stress metabolism (Extended Data Fig. 5c).

**Fig. 4.**
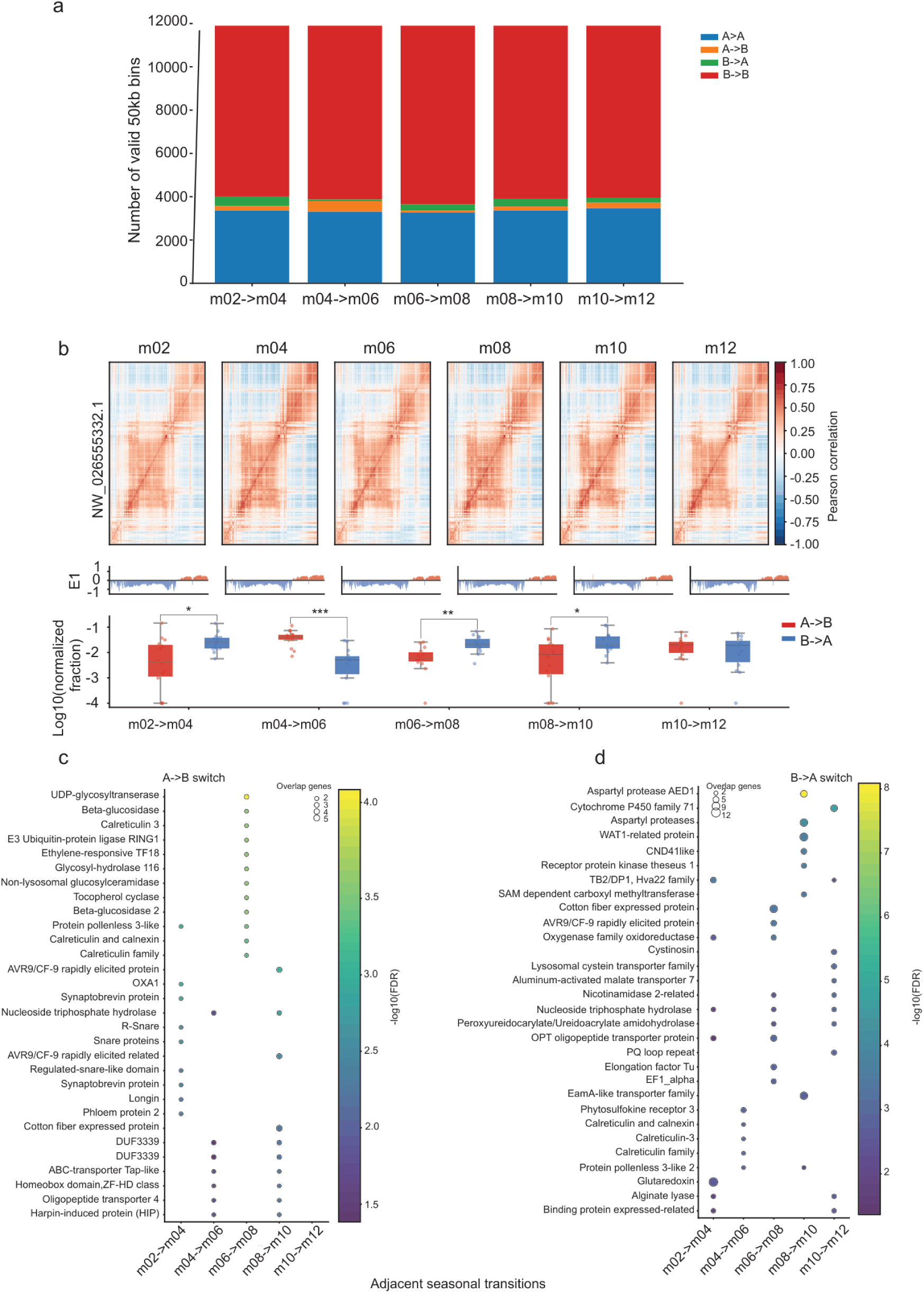
Seasonal A/B compartment switching identifies transition-specific chromatin reprogramming and functional gene classes in *Prosopis cineraria*. **a,** Stacked bar plots show genomic bins assigned to stable A (A→A), stable B (B→B), and switching states (A→B or B→A) across consecutive seasonal transitions. **b,** Representative Pearson correlation maps for scaffold NW_026555332.1 illustrating the spatial organization of compartmental domains. **c,** Functional enrichment of genes associated with A→B switches, including glycosyl hydrolases, ethylene-responsive transcription factors, polygalacturonases, and disease-resistance classes. **d,** Functional enrichment of genes associated with B→A switches, identifying seasonal activation of defense, detoxification, transport, and remodeling pathways.

This transition-specific stress program also extended below ground, where water acquisition is the functional counterpart of heat avoidance. A Prosopis MIZ1-like hydrotropism gene was upregulated during hot seasons and showed concordant H3K4me3 and ATAC-seq gains, indicating active chromatin remodeling at a root water-foraging locus. Moisture-gradient seedling assays showed directional root growth toward wet soil. Quantitative lignin comparison between anatomical regions of curved roots showed stronger deposition in the upper, non-bending region than in the lower bending region (PC1-like contrast: 1.63 versus 1.20, P = 0.0027; PC2-like contrast: 1.27 versus 1.00, P = 0.0473), consistent with localized wall flexibility during hydrotropic curvature rather than a simple gravitropic response. MIZ1 expression increased during the hot season and was positively co-expressed with heat-induced lignin-associated genes, including 4CL, CCR, CAD, laccase, and peroxidase candidates. This coordinated induction is consistent with a heat-associated hydrotropism/lignification candidate module in which root water-seeking behavior is correlated with reinforced lignin deposition and oxidative cell-wall polymerization during peak summer stress (Extended Data Fig. 6). Thus, the same seasonal gating logic that activates wax, transport, detoxification, and stress-remodeling pathways in leaves is associated with root architecture that may improve access to water under peak heat.

### Population and landscape genomics reveal adaptive temporal risk-spreading

Population genomics of 231 individuals resolved seven genetic clusters and non-uniform pairwise FST across the Arabian Peninsula (Fig. 5b,c). XP-CLR scans between cluster 2 and 4 identified peaks near SCARECROW, AGL14, RDH3, LEA/D7, PYL1, ARAD1, PAE9, RCA, CHL, SPL8, F-box, and PLA1 loci (Fig. 5d). Landscape genomics using two methods LFMM2 and RDA identified 7061 SNPs associated with different climate variable to the sampling sites (Supplementary Fig. 8, Supplementary Fig. 9), where the association with the mean temperature of the driest quarter (bio09) identified Bonferroni and FDR-supported SNP-environment signals in ABA perception, hydraulic regulation, ethylene/cytokinin signaling, chloroplast-retrograde communication, PEPC, SCARECROW, TIC32-like, and defense genes. (Fig. 5e) (Data source 1-2). Pollen viability was high in April, moderate in October, and below 5% in June-August (old flowers from the previous season), with significant hot-season variance inflation (Levene W = 90.58, P = 1.08 × 10^-14) and genotype-dependent resilience (Season x Tree likelihood-ratio P = 0.0032), indicating that some genotypes retain a small reproductive margin during peak temperature and is consistent with a temporal risk-spreading by making seeds dormant (Extended Data Fig. 7a) (Supplementary Table 1). Phytohormone profiling revealed a seasonal SA-ABA pattern: salicylic acid peaked in April during the growth/defense phase, whereas ABA doubled during hot months and maximized in August, consistent with stomatal regulation and stress-tolerance induction during peak heat (Extended Data Fig. 7b) (Supplementary Table 2). The qPCR confirmed 10 heat-cluster genes with strong concordance to RNA-seq (overall Pearson r = 0.853, P = 1.52 × 10^-26; gene-average r = 0.857, P = 0.0015). F2 seedlings under simulated cool-heat cycling recapitulated parental heat-cluster directions, consistent with transmitted stress-responsive transcriptional competence rather than proof of fixed transgenerational epigenetic inheritance (Extended Data Fig. 7c; Supplementary Table 3). The stress-memory screen used a staged evidence-integration workflow, reducing 81,383 expressed loci and genomic features to 34,865 memory-like candidates (42.8%) and then to 113 high-confidence priority loci that require RNA persistence, at least one active chromatin layer, and persistence in at least two layers overall. All 113 priority loci showed transcriptional persistence at the m10 post-heat recovery window, and 101 of 113 are also supported by environmental SNPs or XP-CLR sweep evidence; functional classes included ABA/drought signaling, wall-associated receptor kinases, AUX/IAA regulators, amino-acid/polyamine transporters, and RING-type ubiquitin ligases (Extended Data Fig. 8).

**Fig. 5.**
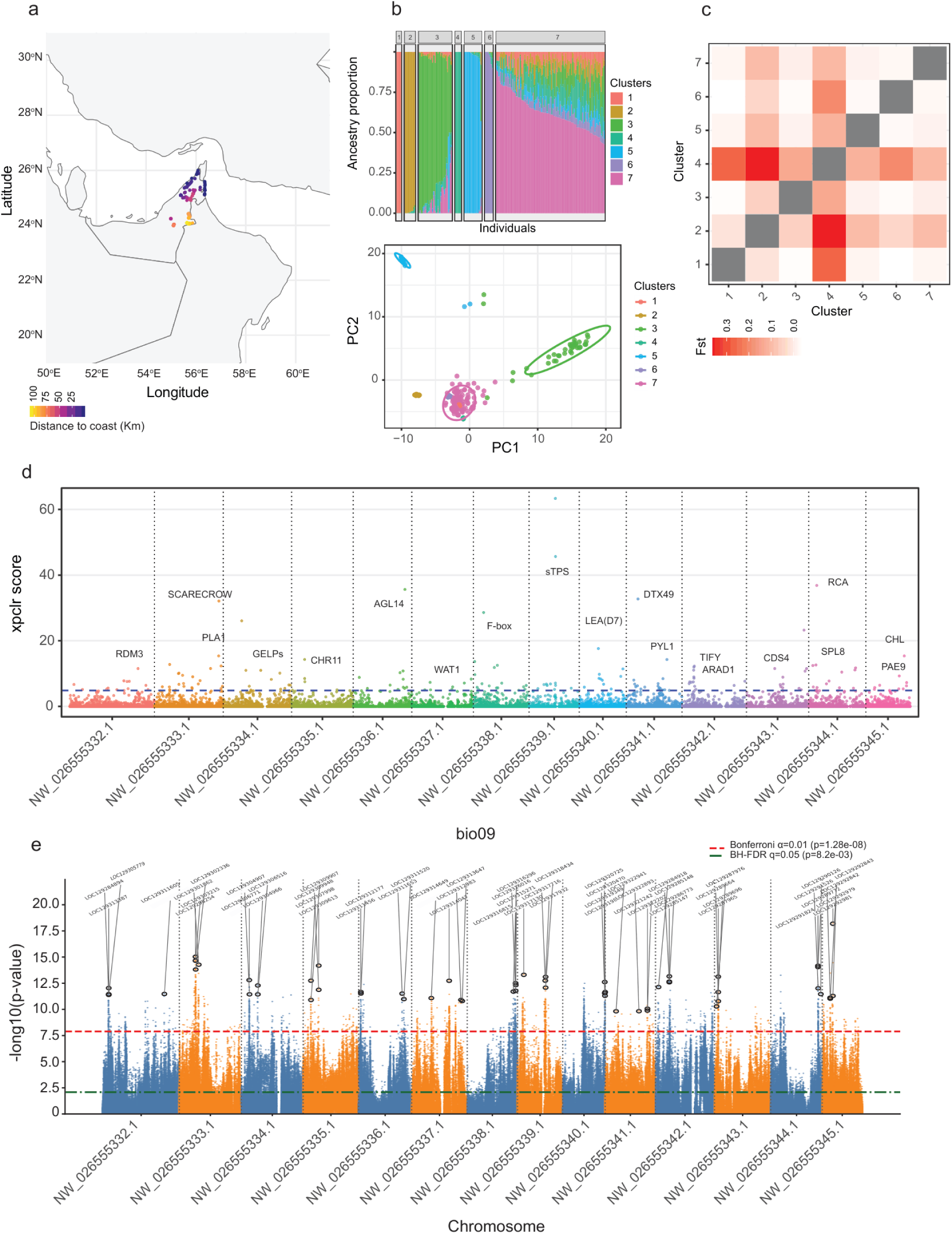
Population structure and environmental genomic signatures of adaptation in *Prosopis cineraria*. **a,** Geographic distribution of 231 sampled individuals across the southern Arabian Peninsula, with points colored by distance to the coast. **b,** Genetic structure inferred from ancestry proportions and PCA revealing seven inferred genetic clusters. **c,** Pairwise genetic differentiation among the seven clusters is shown as an FST heat map. **d,** Genome-wide XP-CLR selective sweep scan across major scaffolds, with labeled peaks marking candidate genes including SCARECROW, RDH3, PYL1, RCA, and SPL8. **e,** LFMM2 Manhattan plot for bio09 (mean temperature of the driest quarter), with Bonferroni and FDR thresholds indicated.

Predicted PEPC stability and Stay-Green regulation support a metabolic-continuity hypothesis. Phylogenetic analysis of PEPC identified a housekeeping PEPC isozyme with ESM2StabP-predicted thermal stability, while alternative splicing analysis highlighted Stay-Green regulation as a candidate contributor to heat-season chlorophyll retention. The Prosopis PEPC housekeeping isozyme clustered with legume homologs and carried 13 specific mutations, including L865, R878, and R885. ESM2StabP predicted a melting temperature of approximately 60.7°C, and residue swaps altered the predicted stability by up to approximately 1.2°C (Fig. 6a). Because these estimates are based on in silico prediction, they should be interpreted as a hypothesis rather than direct evidence of thermostability; protein half-life assays under heat-stress conditions are required to confirm whether this PEPC isoform maintains enhanced stability at the protein level. This isoform is best interpreted as an anaplerotic housekeeping enzyme rather than evidence for a C3-C4 transition. By carboxylating phosphoenolpyruvate to oxaloacetate, a PEPC isoform with predicted heat stability could contribute to MDH-mediated malate interconversion, TCA refilling, redox balance, pH/osmotic regulation, and nitrogen-carbon integration when Rubisco activase becomes vulnerable around 40-45°C; flux measurements are needed to test this pathway. Seasonal malate quantification was compatible with this model but not statistically conclusive, with higher accumulation in the hot June-August group (mean 19.34 ng/µl) than in February-April (13.24 ng/µl) or October-December (14.55 ng/µl); pairwise Wilcoxon tests showed non-significant trends for February-April versus June-August (P = 0.057) and June-August versus October-December (P = 0.057), but not between the two cool-season groups (P = 0.886). Detached-leaf heat-stress assays further showed lower heat-associated chlorophyll loss in P. cineraria than in Medicago sativa across the temperature gradient, supporting chlorophyll stability and recoverable photosynthetic tissue under conditions that bleach a thermosensitive reference species (Extended Data Fig. 9). Proteome-wide predicted-stability enrichments highlighted chaperones, antioxidant enzymes, mitochondrial and TCA metabolism, amino-acid/nitrogen-linked carbon-skeleton metabolism, and core housekeeping proteins. SGR2/SGR1 Stay-Green splicing and expression identified a candidate chlorophyll-retention module that may delay senescence while growth remains restrained (Fig. 6b; Supplementary Fig. 12). Stomatal restriction above approximately 31-32°C reduced the open stomatal fraction without complete shutdown, while chlorophyll content and stem relative humidity were comparatively maintained. CAT, POD, and SOD activities followed non-linear degree-2 polynomial trajectories across the temperature gradient, indicating staged redox adjustment rather than monotonic stress collapse. Together, the PEPC prediction, malate data, stomatal behavior, and redox assays are consistent with a metabolic-continuity hypothesis, but the mechanistic links among PEPC activity, malate flux, and stomatal behavior remain untested (Extended Data Fig. 10a,b; Supplementary Table 7) [64–66].

**Fig. 6.**
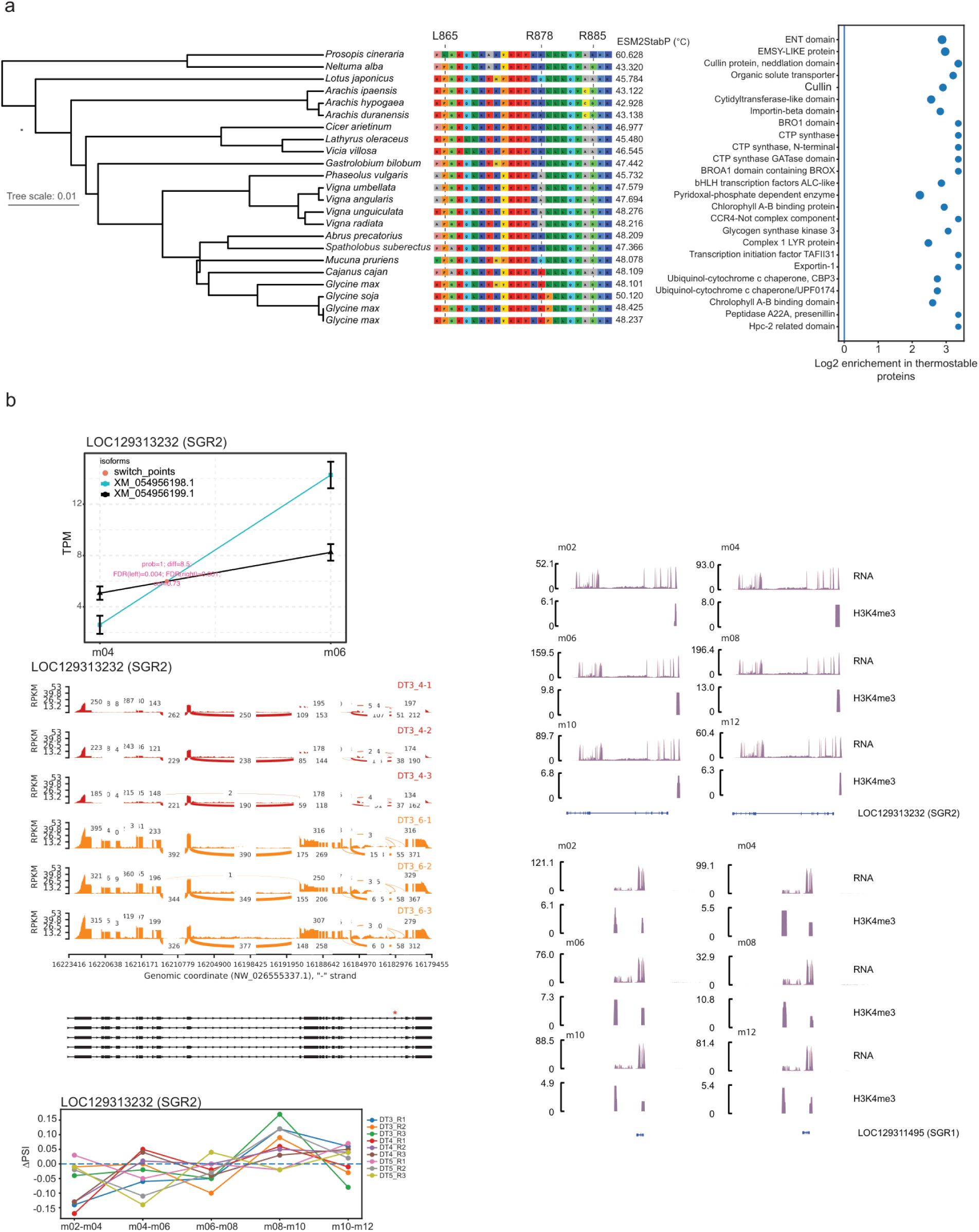
Predicted PEPC stability and stay-green regulation support a heat-season metabolic-continuity hypothesis in Prosopis cineraria. **a**, Predicted protein-stability analysis. (a1) Phylogenetic placement and cropped alignment of Prosopis housekeeping PEPC with legume homologs, highlighting Prosopis residues L865, R878, and R885 and ESM2StabP-predicted melting temperatures. (a2) Log2 enrichment of functional domains among predicted thermostable proteins, implicating core metabolism, transport, chromatin regulation, and mitochondrial/proteostasis functions. b, Seasonal stay-green regulation. (b1) Isoform-level TPM changes for LOC129313232 (SGR2) across m04-m06. (b2) RNA-seq sashimi tracks showing SGR2 exon usage across replicates. (b3) ΔPSI across adjacent seasonal transitions indicating dynamic SGR2 splicing. (b4-b5) Seasonal RNA and H3K4me3 browser tracks for SGR2 and SGR1.

## Discussion

Our results support a model in which Prosopis cineraria survives recurrent desert heat through reversible epigenomic gating rather than constitutive activation of stress pathways. Hot-season boundary weakening, candidate TAD-merge-like patterns, and B-to-A compartment switching were concentrated at stress, transport, wax, detoxification, and metabolic loci, while most chromatin compartments remained stable. This selectivity argues against a generalized heat-induced disruption of chromatin and instead supports regulated seasonal plasticity that expands candidate regulatory contacts at protective loci while avoiding inappropriate genome-wide activation [56,57]. The recovery of insulation in subsequent cool months is central to this interpretation: chromatin reconfiguration appears to be an annual regulatory state rather than cumulative molecular damage.

Active histone marks explain how this architecture becomes transcriptionally productive. H3K4me3 and H3K27ac tracked heat-positive expression more closely than ATAC-seq, indicating that seasonal promoter and enhancer activation, rather than accessibility alone, defines the heat response. Enrichment of NAC, MYB, WRKY, and ERF/AP2 motifs places these chromatin changes within known stress and hormone-response networks. Their convergence with wax genes, antioxidant loci, and MIZ1-linked hydrotropism further connects molecular regulation to organismal traits: leaf surface protection, redox control, and root water foraging.

The MIZ1-linked root phenotype extends this framework from stress perception to mechanical execution. Hydrotropism requires differential growth toward water, and MIZ1 is essential for root hydrotropic bending [55]; the coordinated hot-season induction of MIZ1 with 4CL, CCR, CAD, laccase, and peroxidase candidates suggests that in P. cineraria, water foraging is correlated with spatially controlled secondary-wall reinforcement. Because laccases and class III peroxidases catalyze oxidative lignin polymerization, this module provides a plausible mechanism by which bending tissues remain flexible. At the same time, non-bending regions become lignified and mechanically stabilized during peak heat [62,63].

The promoter-state patterns refine this model. TE-proximal CHH methylation marks cool-season regulatory programs, including flowering and defense, but CHH alone does not establish RNA-directed DNA methylation or repression. Similarly, a low-H3K4me3 hot-season module identifies promoters with reduced active marks, not a demonstrated repressive-histone-mark mechanism. Direct H3K27me3 ChIP-seq, 24-nt small-RNA sequencing, H3K9me2 assays, and allele-specific methylome analyses are required to test those mechanisms. Importantly, recovery of flowering regulators in cool months, together with pollen failure and increased variance during hot months, supports a temporal risk-spreading interpretation while remaining compatible with a simpler heat-sensitivity explanation for old flowers.

Physiology and population genomics strengthen this interpretation: reciprocal seasonal SA and ABA profiles separate cool-season developmental and defense competence from hot-season survival. Landscape-genomic signals in ABA, hydraulic, chloroplast, PEPC, and defense pathways indicate that modules shaped by seasonal chromatin are also targets of adaptation. The overlap between environmental SNP-supported loci and candidate persistence loci suggests convergence between acclimation and evolutionary adaptation. The F2 expression response to simulated seasonal cycles should therefore be interpreted cautiously as heritable stress-responsive competence; plant memory and transgenerational epigenetic frameworks both support this restraint [41,58,59].

The conceptual advance is the linkage of chromatin reversibility to perennial life history. Annual plants can avoid summer heat by completing reproduction, whereas P. cineraria must preserve vegetative and reproductive potential across repeated extremes. Cool-season flowering is therefore not simply phenology, but a risk-spreading pattern: permissive flowering regulators, elevated salicylic acid, and higher pollen viability coincide with periods when reproductive failure is least likely. Conversely, hot months enforce developmental restraint while activating survival investment through ABA accumulation, wax deposition, hydrotropism, stomatal restriction, antioxidant adjustment, and heat-positive chromatin modules. This separation of survival and recovery states is a key value of the multi-omic design; it distinguishes a coordinated seasonal program from a generic drought response [60].

Several limitations define the next experiments. Low-H3K4me3 hot-season repression requires direct H3K27me3 ChIP-seq across the same six seasons, while TE-proximal CHH methylation should be tested with 24-nt small-RNA profiles, H3K9me2, and strand-specific methylomes. The PEPC model needs biochemical validation via enzyme activity (half-life at 60 °C) and TCA-flux measurements, including comparisons of native and residue-swapped variants. A future Stay-Green transient-expression assay should test whether SGR2-associated isoforms alter SGR1-linked chlorophyll degradation. Finally, F2 seedlings should be profiled epigenomically to separate inherited chromatin memory from genetically encoded inducibility.

Despite these caveats, the integrated evidence has clear conservation implications. *P. cineraria* populations harbor spatially structured genetic diversity and adaptive variants associated with the driest-quarter temperature. However, persistence also depends on plastic regulatory systems that allow individuals to traverse unfavorable seasons. Conservation strategies based only on neutral diversity would miss this functional dimension. Seed sourcing, restoration, and assisted adaptation should therefore consider both climate-associated allelic variation and seasonal epigenomic responsiveness. The same logic applies to crop improvement: heat tolerance should be selected not only for acute heat-shock resistance, but also for regulatory timing, recovery capacity, and metabolic buffering.

This framing also guards against over-interpretation. CHH methylation can coexist with active genes and therefore cannot alone define repression. ATAC-seq may mark competence without predicting RNA output. The predicted PEPC stability result does not imply C4 evolution; at most, it supports a testable hypothesis that a housekeeping PEPC contributes to anaplerosis and malate production under heat. Pollen collapse alone indicates heat sensitivity, but together with cool-season flowering, genotype-dependent variance, and flowering-gene recovery, it supports a temporal risk-spreading interpretation. The central claim is therefore not that P. cineraria possesses a single exceptional trait, but that multiple reversible systems are coordinated seasonally across chromatin topology, promoter marks, methylation, hormones, roots, reproduction, and predicted protein stability.

Predicted PEPC stability and Stay-Green regulation provide a candidate biochemical framework for reversible growth restraint. When Rubisco activase becomes heat-labile, a predicted heat-stable housekeeping PEPC could help maintain oxaloacetate and malate production for TCA-cycle carbon supply, osmotic balance, and repair metabolism, but this remains to be tested directly. In parallel, SGR2-associated chlorophyll retention may preserve green tissue, allowing photosynthetic capacity to be reactivated after heat. We therefore propose a growth-off, metabolic-core-on model: chromatin gates seasonal transcription; hormone profiles align with allocation shifts; roots seek water; candidate protein stability may maintain core metabolism; and Stay-Green may delay irreversible senescence. This framework highlights desert perennials as useful models for understanding adaptation to warming extremes.

Beyond conservation, these findings suggest translational-genomics routes for heat-sensitive legumes. The predicted heat-stable housekeeping PEPC identified here should be viewed not as a photosynthetic C4-like shift, but as a candidate module for preserving anaplerotic malate production, pH balance, and carbon-skeleton supply when Calvin-cycle flux becomes heat constrained [28,31]. Introgression, allele mining, genome editing, or synthetic reconstruction of this PEPC-associated regulatory circuit could be tested for metabolic continuity under episodic heat. Stay-Green-mediated delay of chlorophyll catabolism offers a complementary candidate strategy to preserve recoverable photosynthetic tissue and avoid premature senescence [32]. These ideas align with recent calls to mine wild plant adaptations for crop heat tolerance. However, they must be tested for yield, phenology, and stress-response persistence trade-offs under realistic fluctuating heat and drought regimes [61].

A parallel but more speculative opportunity concerns flowering recovery. The cool-season flowering module suggests that H3K4me3-linked permissive chromatin may help preserve reproductive competence until temperatures become favorable, rather than forcing flowering during lethal heat. In sensitive legumes, this should be framed as a testable route to identify homologous flowering circuits whose chromatin responsiveness can be selected, primed, or edited to improve post-heat floral recovery and extend reproductive windows [19,37]. This caution is consistent with the view that wild stress-adapted plants provide valuable breeding targets. However, heat tolerance is often polygenic, redundant, and constrained by growth-survival trade-offs [61].

Overall, the study defines desert resilience as seasonal allocation rather than simple tolerance. During cool months, *P. cineraria* initiates flowering, immune competence, and developmental recovery; during hot months, it redirects resources toward water seeking, surface protection, redox control, hormonal restraint, and preservation of the metabolic core; after heat, chromatin boundaries and cool-season programs return. This cyclic plasticity is likely to be essential for long-lived perennials facing a harsh environment, because survival depends not only on withstanding maximum temperature, but also on preserving the capacity to recover.

## Supporting information

Extended data figures mentioned throughout the manuscript

## Data availability

All raw data generated in this study have been deposited in NCBI under the accession number PRJNA******. Datasets used in this study have been compiled in GitHub (through IGV-web) at https://github.com/KCGEB-Admin/Ghaf-Hub/tree/main/Ghaf_multiomics_browser.

## Code availability

Analysis scripts used in this study can be found on GitHub at https://github.com/KCGEB-Admin/Ghaf-Hub/tree/main/Prosopis_manuscript_bioinformatics_scripts.

## Acknowledgements

The authors would like to thank and acknowledge the United Arab Emirates University (UAEU; Al Ain, UAE) and the Khalifa Center for Genetic Engineering and Biotechnology (KCGEB; Al Ain, UAE) for providing laboratory facilities and additional resources.

## Author contributions

Conceptualization and study design: KMH, KMAA; Methodology (Sample collection & processing): BD, RM, FTP, DK, IS, TR, MP, KMH; Analysis (Data curation): BD, KMH, and KMAA; Funding acquisition: KMAA; Writing—original draft: BD and KMH; Writing review and editing: BD, KMH, and KMAA.

## Competing interests

The authors declare no competing interests.

## Notes

### Competing Interest Statement

The authors have declared no competing interest.

### Summary of Updates

We would like to withhold the NCBI accession number for data availability until the final peer-review submission to prevent data leakage and being scooped.

