## Supplementary material for "Reversible chromatin remodeling enables *Prosopis cineraria* survival under recurrent heat extremes": Supplimentary information

**Table of Contents**

Figure S1. Hot seasons show a modest increase in the fraction of large TADs.

Figure S2. Time-series clustering of seasonally dynamic gene-expression profiles.

Figure S2. Seasonal chromatin accessibility and H3K27ac profiles across stress-responsive gene clusters.

Figure S4. Active A compartments coincide with H3K4me3-enriched chromatin (e.

Figure S5. Heat-season induction of VIL2–PRC2/chromatin regulators supports an H3K27me3-linked FLC repression module.

Figure S6. Seasonal flowering-gene activation contrasts with heat-associated FLC/FLX repressor induction.

Figure S7. Landscape-genomic scans identify shared and method-specific climate-associated loci.

Figure S8. Genome-wide landscape-genomic associations with climatic and edaphic variables.

Table S1. Summary of aniline blue-based pollen viability across seasonal sampling points.

Table S2. Seasonal hormone summary across sampling months.

Table S3. Primer sequences for selected candidate genes used for qPCR validation.

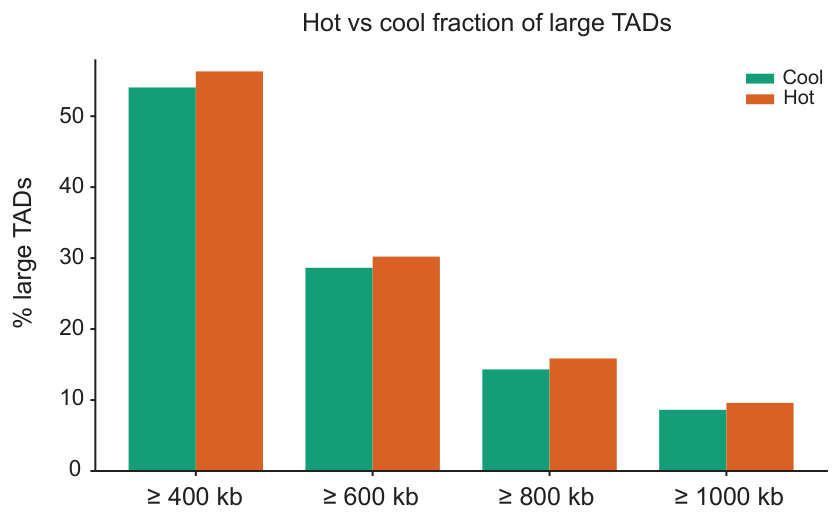

**Figure S1. Hot seasons show a modest increase in the fraction of large TADs.** Bar plot comparing the proportion of topologically associating domains (TADs) classified as large across cool and hot seasons using increasing size thresholds (≥400, ≥600, ≥800 and ≥1000 kb). Hot-season samples show consistently higher fractions of large TADs across all thresholds, supporting a heat-associated shift toward larger chromatin domains and possible TAD merging during peak thermal stress.

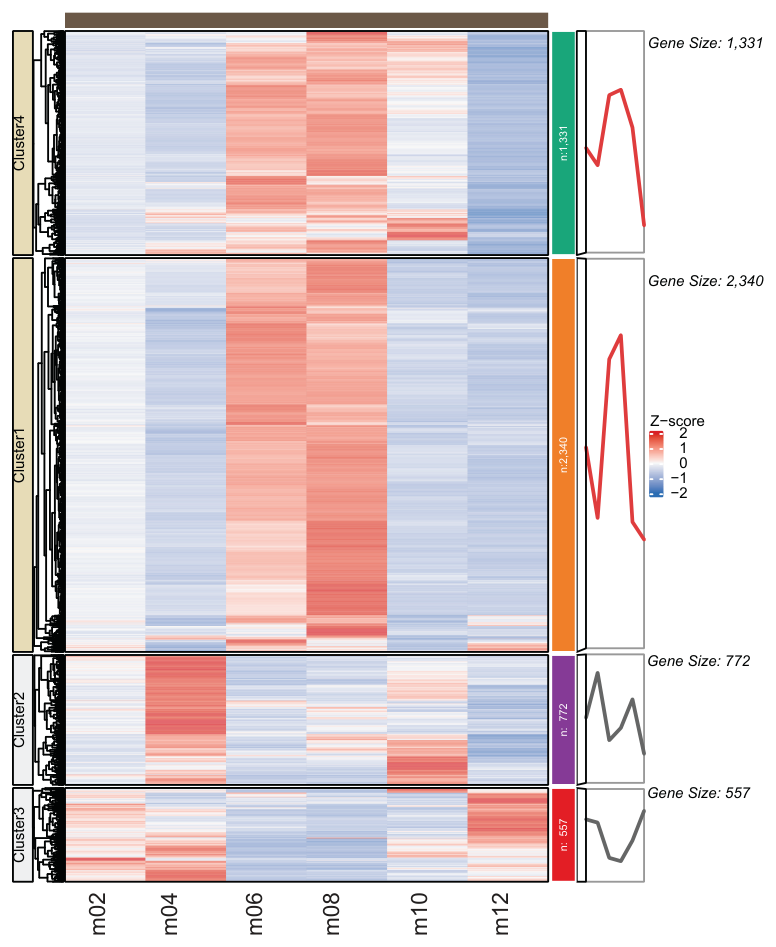

**Figure S2. Time-series clustering of seasonally dynamic gene-expression profiles.** Heat map showing z-score-normalised expression profiles of seasonally variable genes across six sampling months: February (m02), April (m04), June (m06), August (m08), October (m10) and December (m12). Genes were grouped into four temporal clusters according to similarity in seasonal expression dynamics and visualised using the ComplexHeatmap R package (Gu et al., 2016). Cluster4 contained 1,331 genes and Cluster1 contained 2,340 genes, both showing heat-associated induction with elevated expression during the peak hot months, particularly June–August. Cluster2 contained 772 genes and showed a transition-season pattern with stronger expression around April and October. Cluster3 contained 557 genes and showed a cool-season-enriched profile, with higher expression in February/December and reduced expression during the hot-season interval. Rows represent genes and columns represent seasonal time points; colours indicate relative expression z-scores, with red denoting higher and blue denoting lower expression. Line plots on the right summarize the average temporal profile for each cluster.

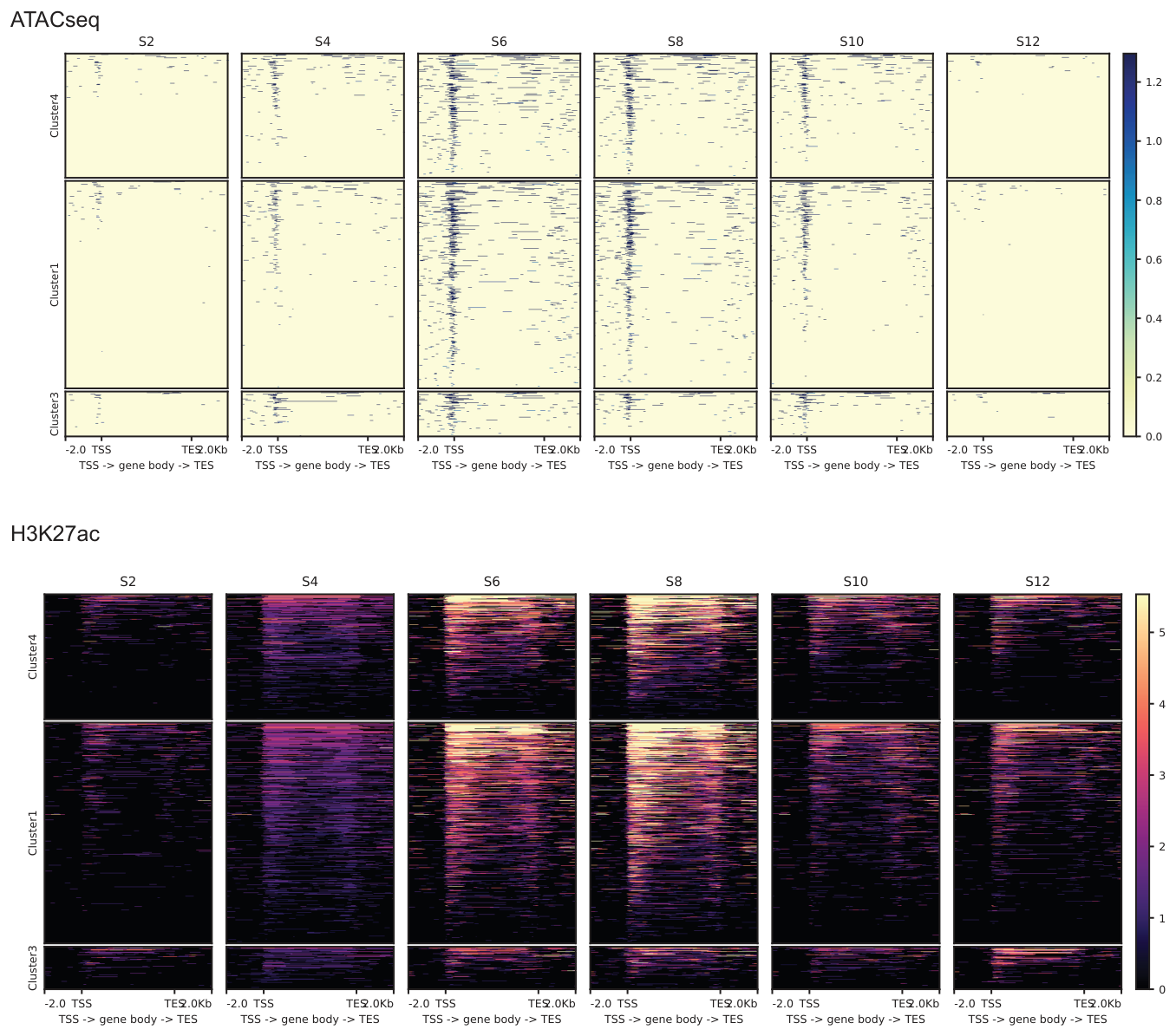

**Figure S3. Seasonal chromatin accessibility and H3K27ac profiles across stress-responsive gene clusters.** Heat maps showing ATAC-seq and H3K27ac signal across gene bodies and ±2 kb flanking regions from TSS to TES for three seasonal gene clusters across S2, S4, S6, S8, S10 and S12. ATAC-seq accessibility and H3K27ac enrichment increase most prominently during hot seasons S6–S8, particularly in heat-responsive clusters, indicating coordinated seasonal activation of open chromatin and enhancer-associated acetylation during peak thermal stress.

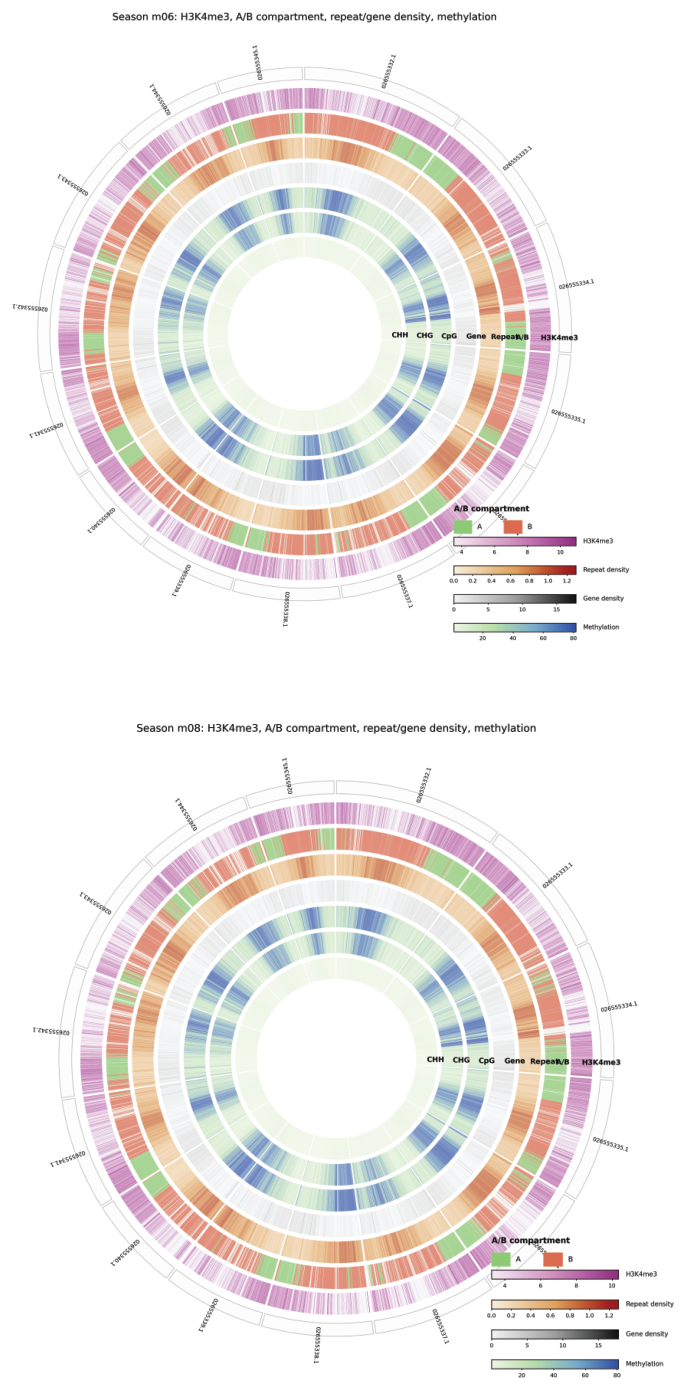

**Figure S4. Active A compartments coincide with H3K4me3-enriched chromatin (e.g., hot seasons) Circos plots for the hot seasons m06 and m08 showing genome-wide relationships among A/B compartment status, H3K4me3 signal, repeat density, gene density and CG, CHG and CHH methylation across the major scaﬀolds.** A-compartment regions show broad overlap with elevated H3K4me3 and gene-rich chromatin, whereas B-compartment regions are generally more repeat- and methylation-associated, supporting a positive relationship between active 3D genome compartmentalization and H3K4me3-marked transcriptionally permissive chromatin during peak seasonal heat.

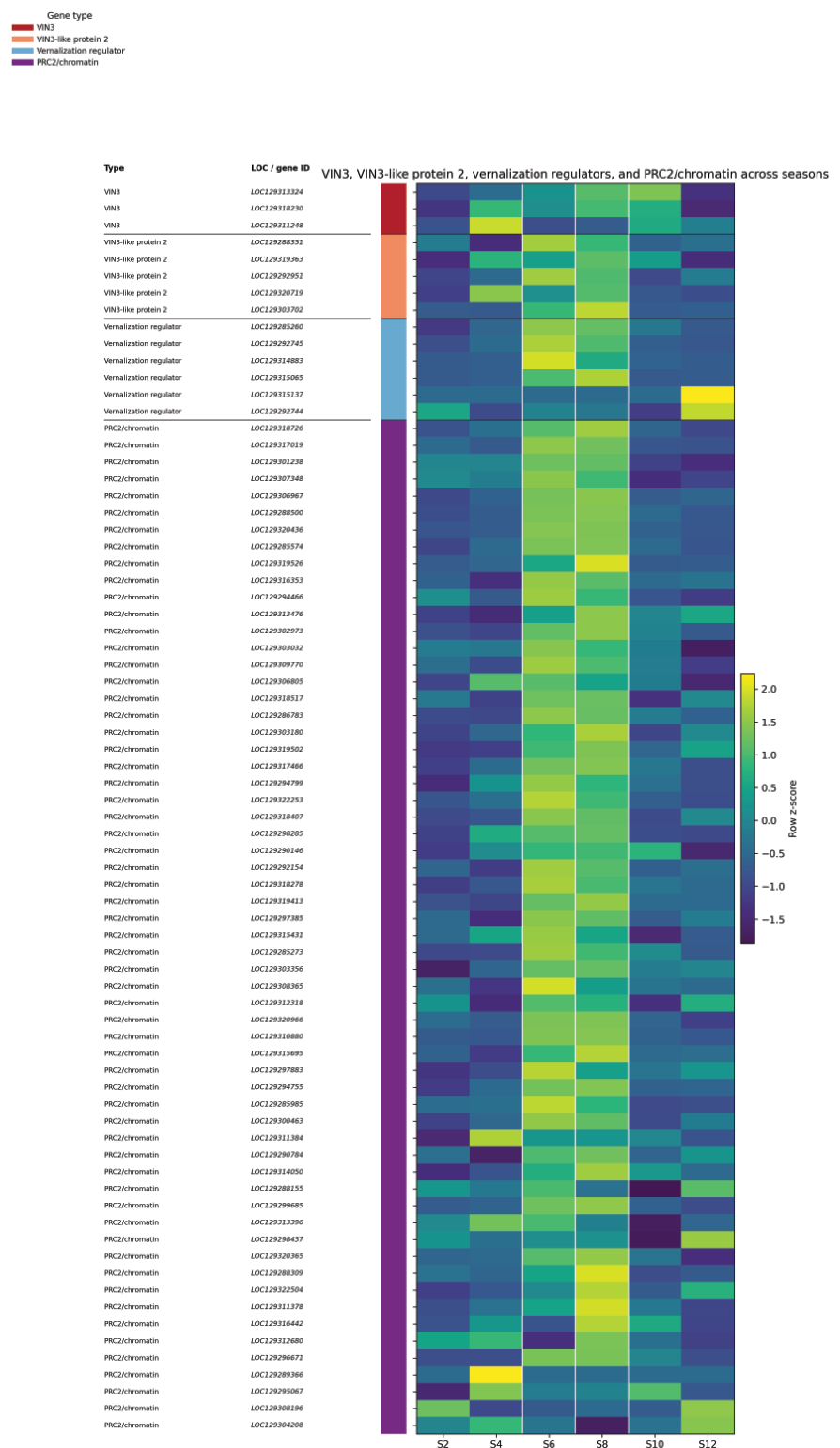

**Figure S5. Heat-season induction of VIL2–PRC2/chromatin regulators supports an H3K27me3-linked FLC repression module.** Heat map showing row-scaled expression of VIN3, VIN3-like protein 2, vernalization regulators and PRC2/chromatin-associated genes across seasons S2, S4, S6, S8, S10 and S12. Unlike the cold-induced VIN3, the constitutively expressed VIN3-LIKE PROTEIN 2 (VIL2) is coexpressed with PRC2 components to support H3K27me3-mediated epigenetic repression at specific photoperiodic target loci. The coordinated induction of VIL2, vernalization-associated regulators and PRC2/chromatin genes during the hot seasons S6–S8 is consistent with activation of a PRC2/H3K27me3-like repression module that may contribute to heat-associated repression of FLC-related flowering regulators and seasonal growth control.

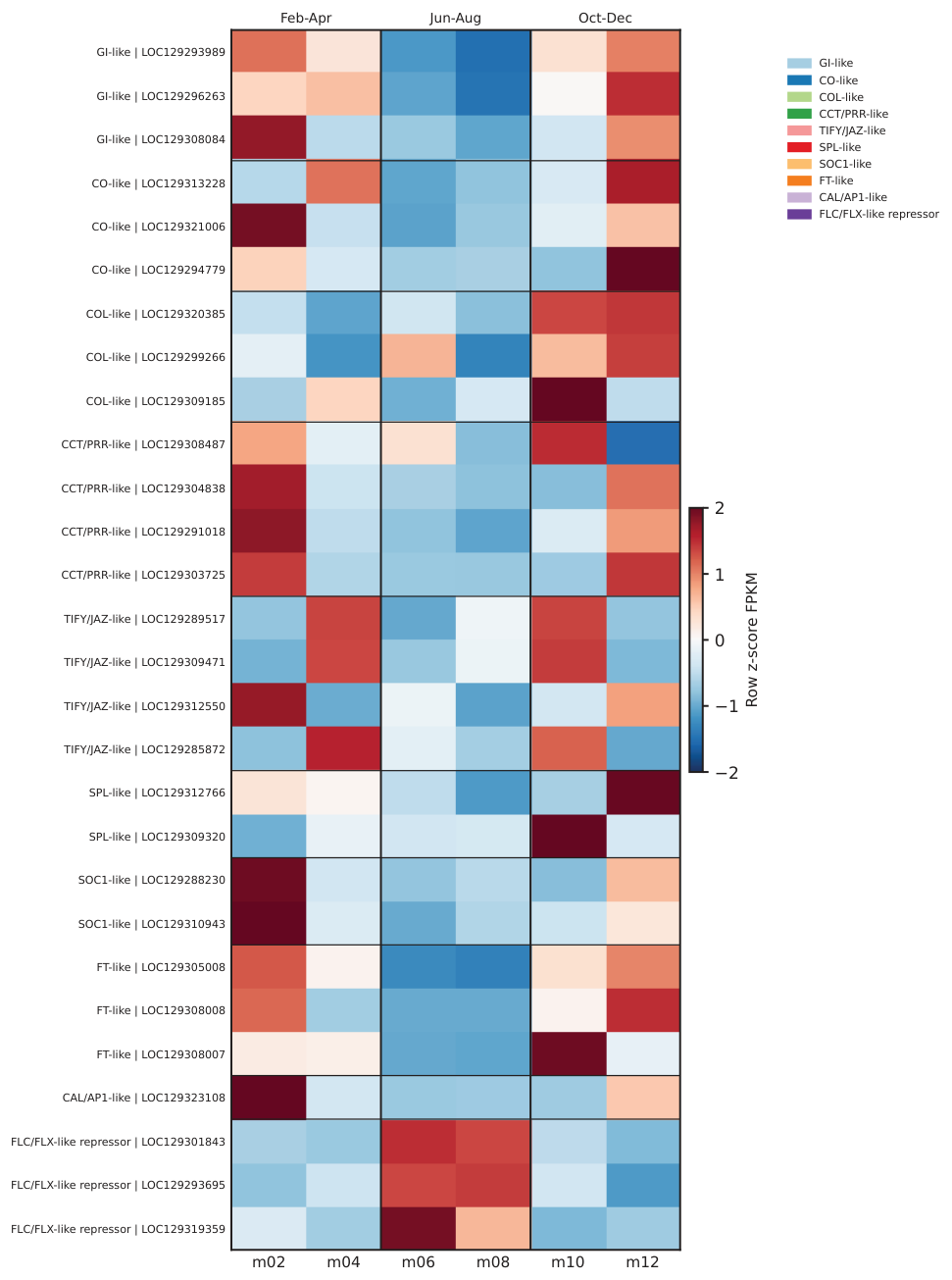

**Figure S6. Seasonal flowering-gene activation contrasts with heat-associated FLC/FLX repressor induction.** Heat map showing row-scaled FPKM expression of flowering-time and photoperiod-associated genes across six seasons (m02, m04, m06, m08, m10 and m12). GI-, CO/COL-, CCT/PRR-, TIFY/JAZ-, SPL-, SOC1-, FT- and CAL/AP1-like genes show stronger activation during cool flowering-permissive periods, particularly Feb–Apr and Oct–Dec. In contrast, FLC/FLX-like repressors are preferentially induced during the hot seasons m06–m08, consistent with heat-associated repression of flowering output and seasonal suppression of reproductive transition under peak thermal stress.

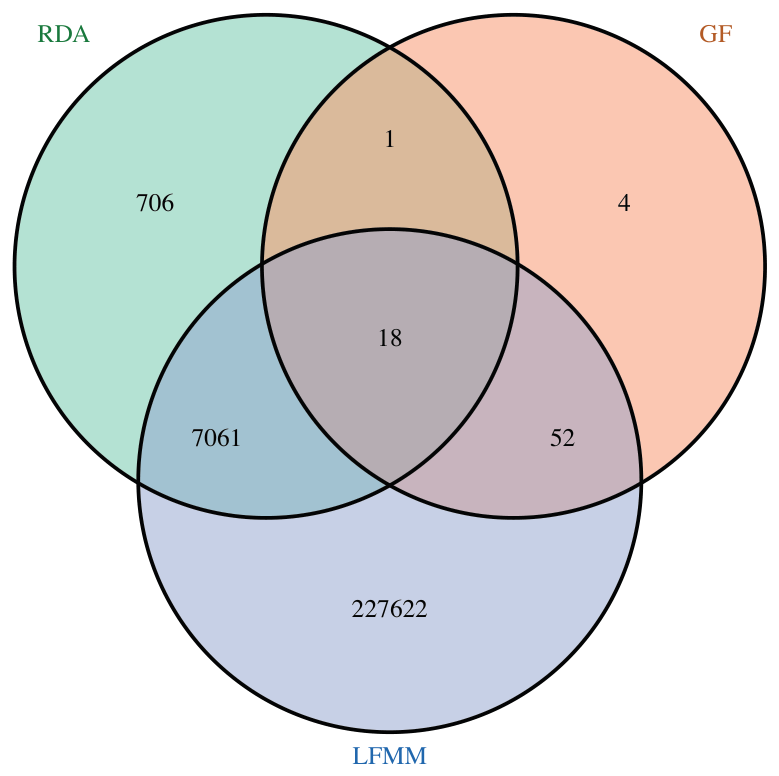

**Figure S7. Landscape-genomic scans identify shared and method-specific climate-associated loci.** Venn diagram showing overlap among candidate loci detected by three landscape-genomic approaches: redundancy analysis (RDA), gradient forest (GF) and latent factor mixed models (LFMM). Most candidates were method-specific, especially for LFMM, whereas the large RDA–LFMM intersection identified 7,061 shared loci, indicating broad concordance between multivariate and latent-factor genotype–environment association tests. A smaller set of 18 loci was recovered by all three approaches, representing high-confidence environment-associated candidates potentially involved in local adaptation to heat and aridity.

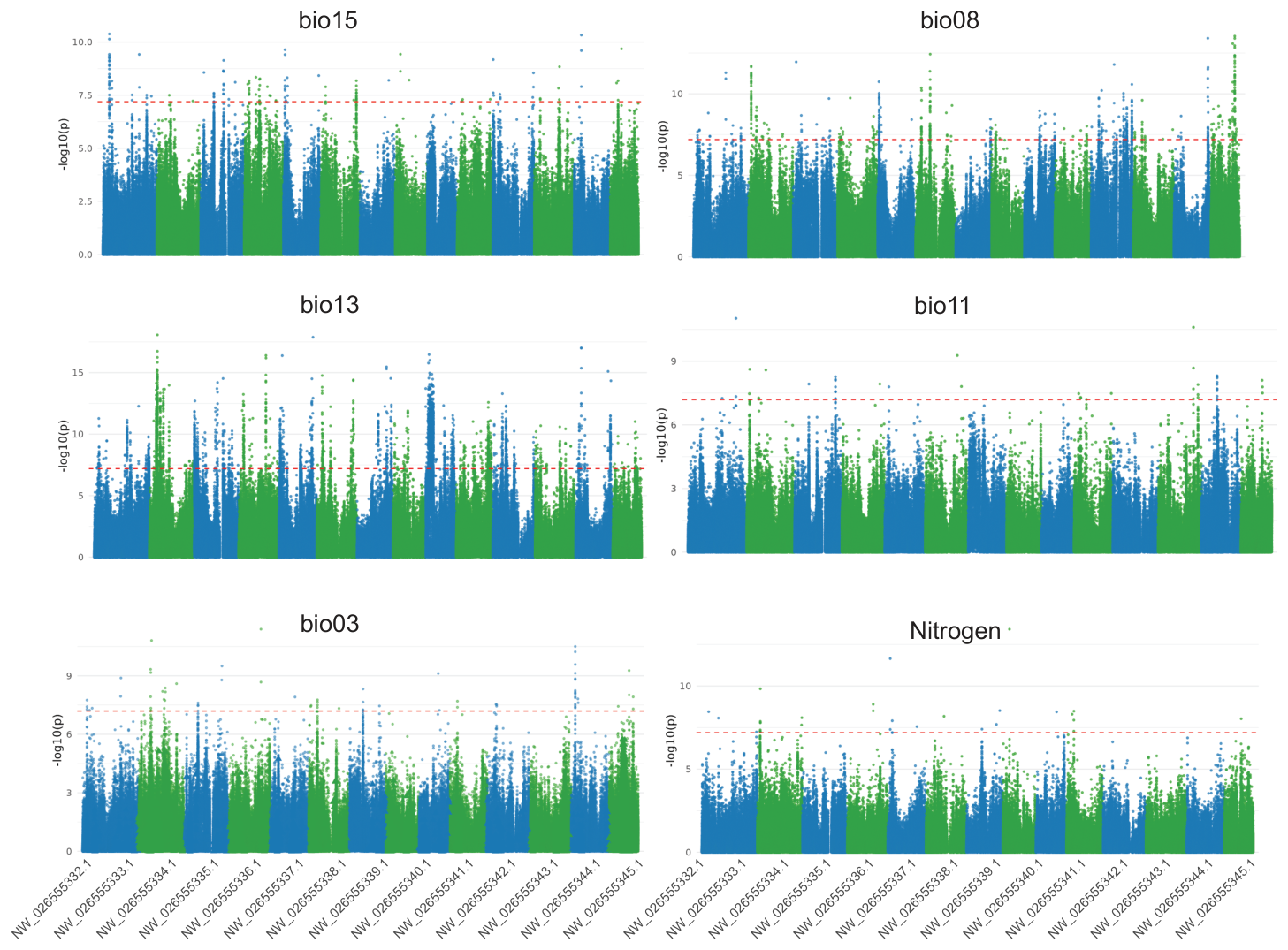

**Figure S8. Genome-wide landscape-genomic associations with climatic and edaphic variables.** Manhattan plots showing SNP–environment association signals across the 14 major scaﬀolds for six environmental predictors: bio15, bio08, bio13, bio11, bio03 and nitrogen. Alternating scaﬀold colors indicate chromosomal positions, and the dashed horizontal line marks the candidate significance threshold. Peaks above the threshold identify genomic regions associated with precipitation seasonality, temperature and soil nitrogen variation, supporting a polygenic landscape-genomic basis for local adaptation to heterogeneous desert environments.

**Table S1. Summary of aniline blue-based pollen viability across seasonal sampling points.** Values are summarized by season as means and standard errors (SE) across scored flowers; 500 pollen grains were counted per flower in the original scoring table.

| **Season** | **No. flowers** | **Total pollen counted** | **Mean viable pollen count** | **SE viable pollen count** | **Mean nonviable pollen count** | **SE nonviable pollen count** | **Mean viability (%)** | **SE viability (%)** |
| --- | --- | --- | --- | --- | --- | --- | --- | --- |
| m04 | 20 | 10000 | 422.60 | 5.65 | 77.40 | 5.65 | 84.52 | 1.13 |
| m06 | 20 | 10000 | 18.60 | 4.61 | 481.40 | 4.61 | 3.72 | 0.92 |
| m08 | 20 | 10000 | 19.30 | 5.57 | 480.70 | 5.57 | 3.85 | 1.11 |
| m10 | 20 | 10000 | 303.25 | 3.91 | 196.75 | 3.91 | 60.64 | 0.78 |

**Table S2. Seasonal hormone summary across sampling months.** Mean, standard deviation and standard error values are reported for each hormone measured across seasonal sampling months.

| **Month** | **Season** | **Hormone** | **Mean** | **SD** | **n** | **SEM** |
| --- | --- | --- | --- | --- | --- | --- |
| April | April | ACC (Ethylene) | 1,710.42 | 1,992.26 | 3 | 1,150.23 |
| April | April | Abscisic acid | 759.85 | 494.43 | 3 | 285.46 |
| April | April | Gibberellic acid | 281.34 | 26.00 | 3 | 15.01 |
| April | April | Salicylic acid | 36,072.71 | 16,379.30 | 3 | 9,456.59 |
| Aug | Aug | ACC (Ethylene) | 950.58 | 827.37 | 3 | 477.68 |
| Aug | Aug | Abscisic acid | 5,315.80 | 1,534.87 | 3 | 886.16 |
| Aug | Aug | Gibberellic acid | 457.12 | 68.88 | 3 | 39.77 |
| Aug | Aug | Salicylic acid | 5,033.59 | 5,426.88 | 3 | 3,133.21 |
| Dec | Dec | ACC (Ethylene) | 398.13 | 192.97 | 3 | 111.41 |
| Dec | Dec | Abscisic acid | 4,776.03 | 1,449.57 | 3 | 836.91 |
| Dec | Dec | Gibberellic acid | 491.95 | 56.48 | 3 | 32.61 |
| Dec | Dec | Salicylic acid | 20,685.84 | 5,353.26 | 3 | 3,090.71 |
| Feb | Feb | ACC (Ethylene) | 16,367.08 | 25,777.61 | 3 | 14,882.71 |
| Feb | Feb | Abscisic acid | 2,487.85 | 663.52 | 3 | 383.08 |
| Feb | Feb | Gibberellic acid | 281.99 | 82.27 | 3 | 47.50 |
| Feb | Feb | Salicylic acid | 25,152.07 | 5,174.45 | 3 | 2,987.47 |
| June | June | ACC (Ethylene) | 2,199.01 | 1,171.07 | 3 | 676.12 |
| June | June | Abscisic acid | 4,583.23 | 433.45 | 3 | 250.25 |
| June | June | Gibberellic acid | 487.15 | 15.22 | 3 | 8.78 |
| June | June | Salicylic acid | 12,432.17 | 1,504.12 | 3 | 868.40 |
| Oct | Oct | ACC (Ethylene) | 1,367.12 | 1,077.12 | 3 | 621.87 |
| Oct | Oct | Abscisic acid | 1,485.14 | 1,028.15 | 3 | 593.60 |
| Oct | Oct | Gibberellic acid | 618.99 | 150.70 | 3 | 87.01 |
| Oct | Oct | Salicylic acid | 13,494.59 | 17,616.42 | 3 | 10,170.85 |

**Table S3. Primer sequences for selected candidate genes used for qPCR validation.** The table lists the ten candidate genes highlighted in the gene panel, with primer names, forward and reverse oligonucleotide sequences (5′–3′), and primer length calculated directly from each sequence.

| **Gene** | **Forward primer** | **Forward sequence (5′–3′)** | **F length (nt)** | **Reverse primer** | **Reverse sequence (5′–3′)** | **R length (nt)** |
| --- | --- | --- | --- | --- | --- | --- |
| LTP2 | PC-LTP2-1-F | TGCTTTGTGTGCACTGTTGGT | 21 | PC-LTP2-1-R | GCACTGGGCTGCATGTTACT | 20 |
| FLA1 | PC-FLA1-1-F | GTGATGGTGGTCTTGCCGTT | 20 | PC-FLA1-1-R | GCCAAGCATCCGGAGTTTTCT | 21 |
| ATXR | PC-ATXR-F | CCATGGTCGTGAGAAGTCGC | 20 | PC-ATXR-R | TGTCTTGTGGGGACTCAGCA | 20 |
| ABC transporter | ABC transporter-F | GCACTGTAGGGCTGGATAG | 19 | ABC transporter-R | GGGTTCTCCATGCCTAA | 17 |
| GLK1 | GLK1-F | TGACACTCGAGATAGATAG | 19 | GLK1-R | CTCCACCTGGCTGATATT | 18 |
| PEPC3 | PEPC3-F | CCTCAGCTTCAATGGTAGTC | 20 | PEPC3-R | CCGACTTCGCGATTCATACA | 20 |
| HOP | HOP-F | GTGAAATCACCGATGAGA | 18 | HOP-R | CAACTCGTCACTCTCTTTC | 19 |
| ENOD | ENOD-F | GGACAAGCTGAATCTGTACC | 20 | ENOD-R | CCAGCATGGCTTCTTGAT | 18 |
| HSP17.9 | HSP17.9-F | GTAGCAACACAGAGAGAAG | 19 | HSP17.9-R | CCTTCTGTCAAGTCCATCAT | 20 |
| SRH3 | SRH3-F | CGACGAGCCTTTGGCTAA | 18 | SRH3-R | GGCTGAATCTCCGAACGA | 18 |
