## Supplementary figures and images for "Reversible chromatin remodeling enables *Prosopis cineraria* survival under recurrent heat extremes"

### Extended data figures mentioned throughout the manuscript

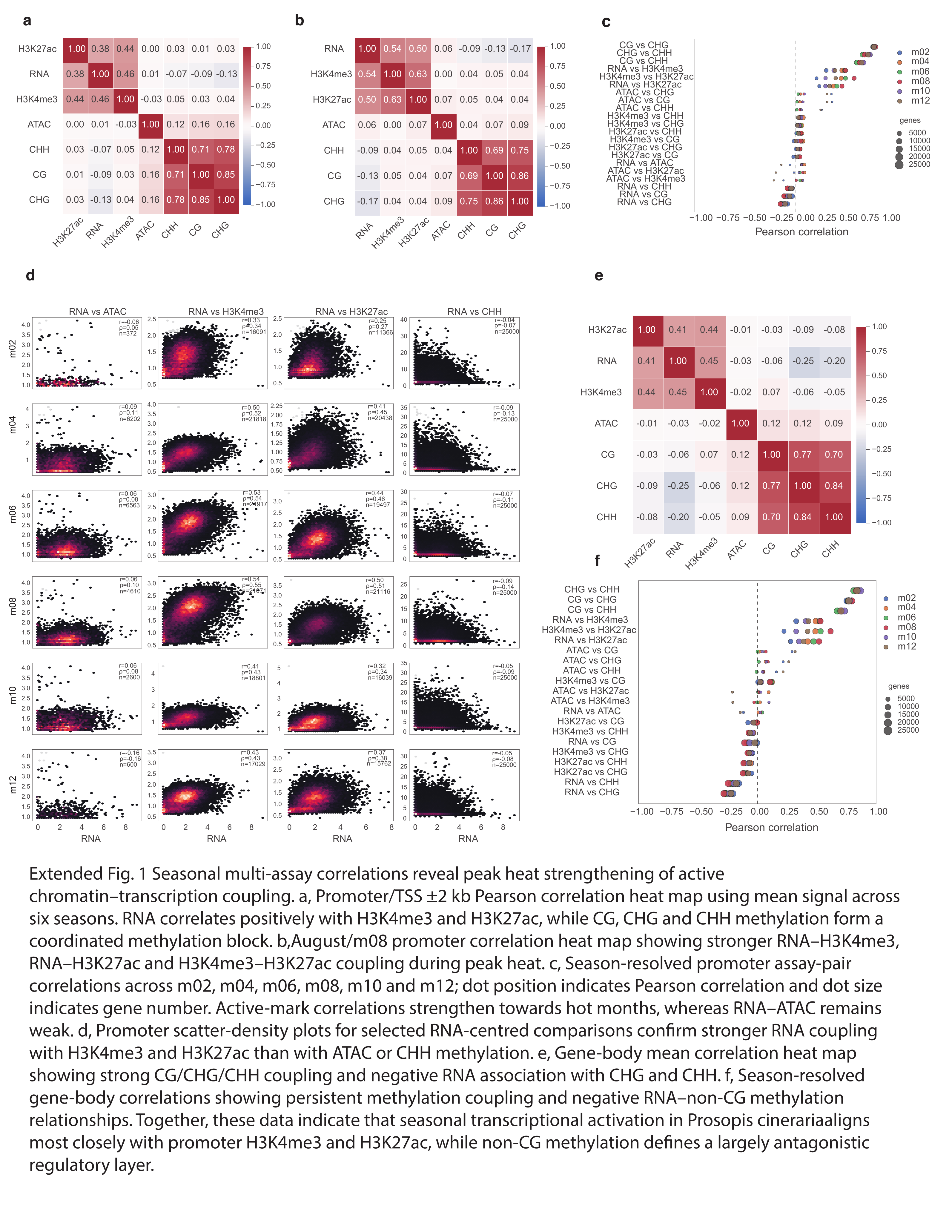

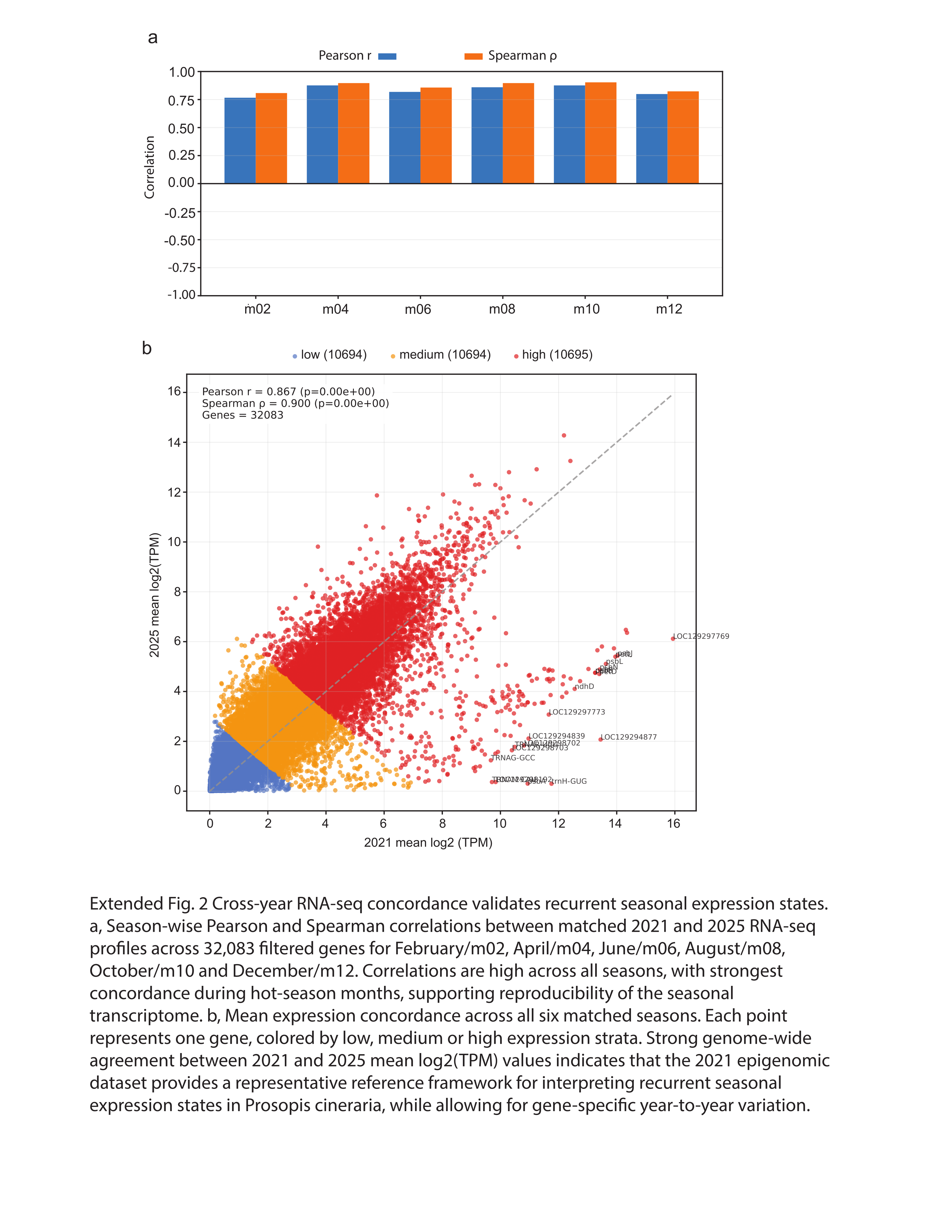

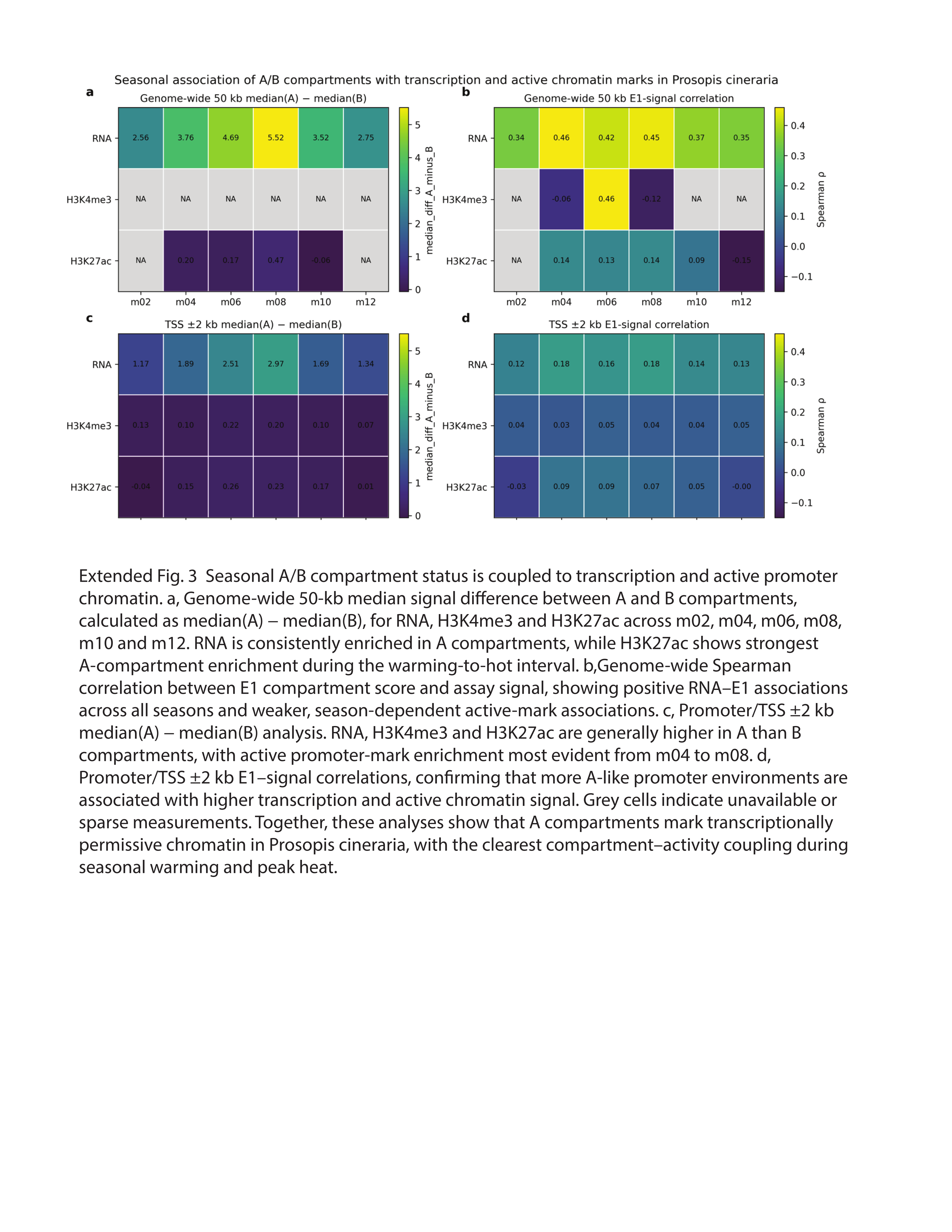

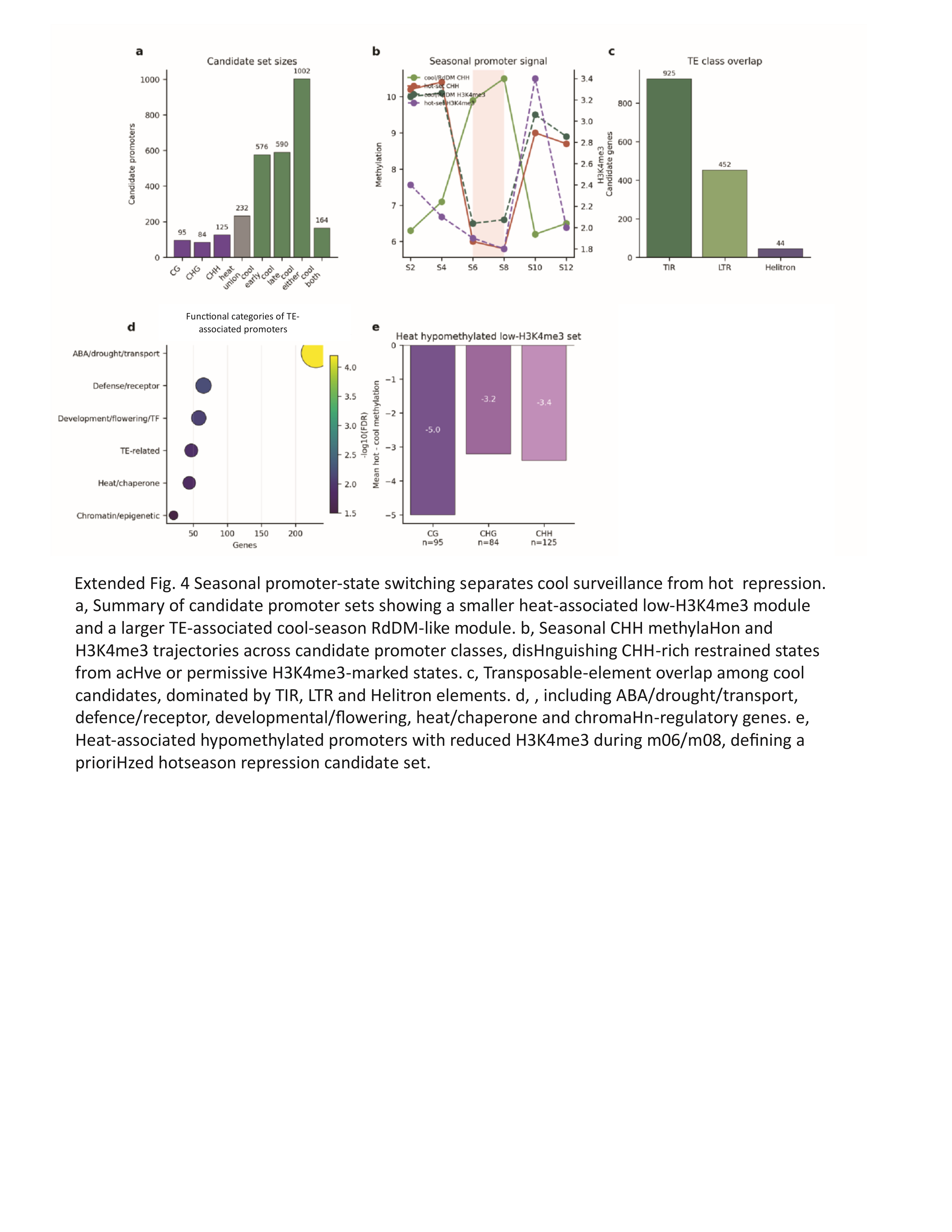

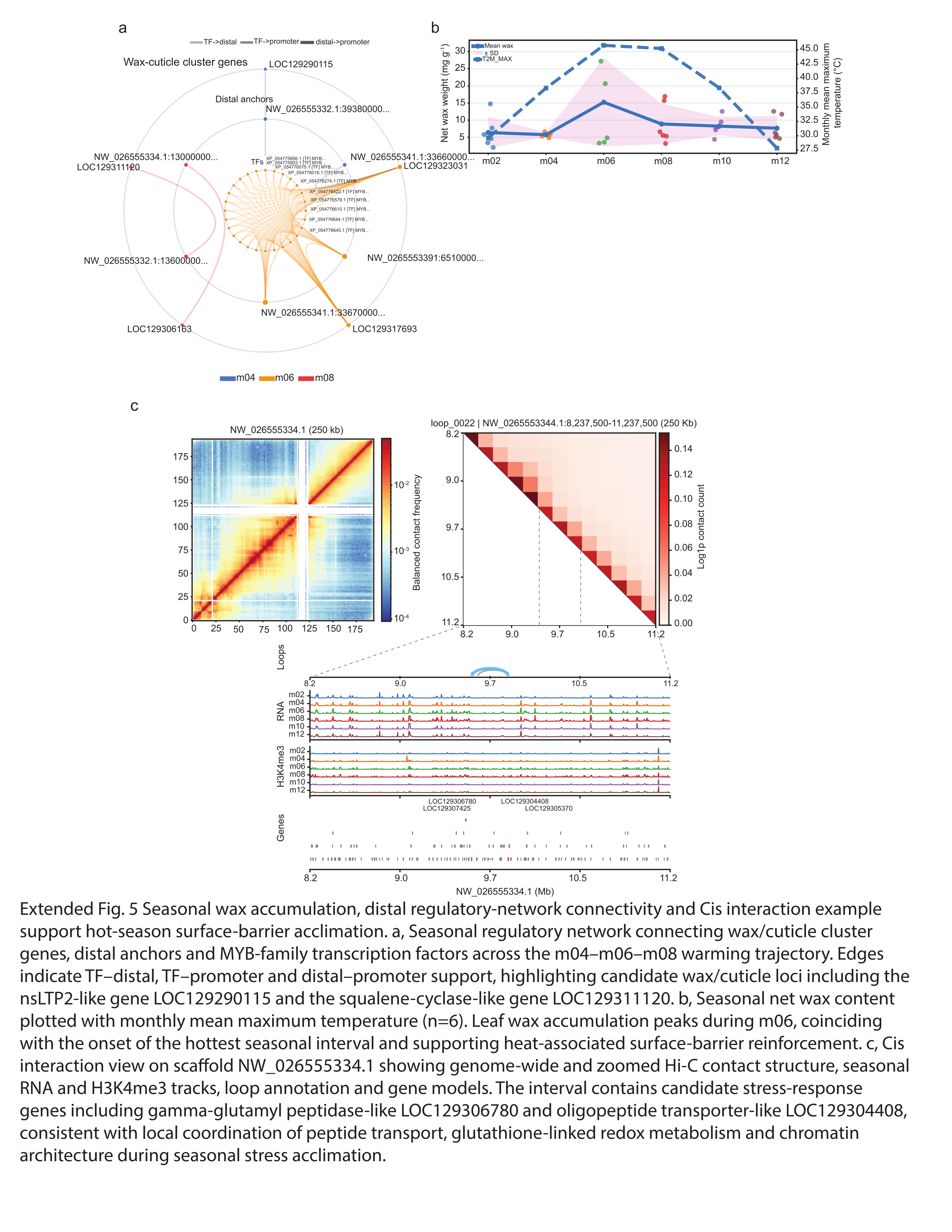

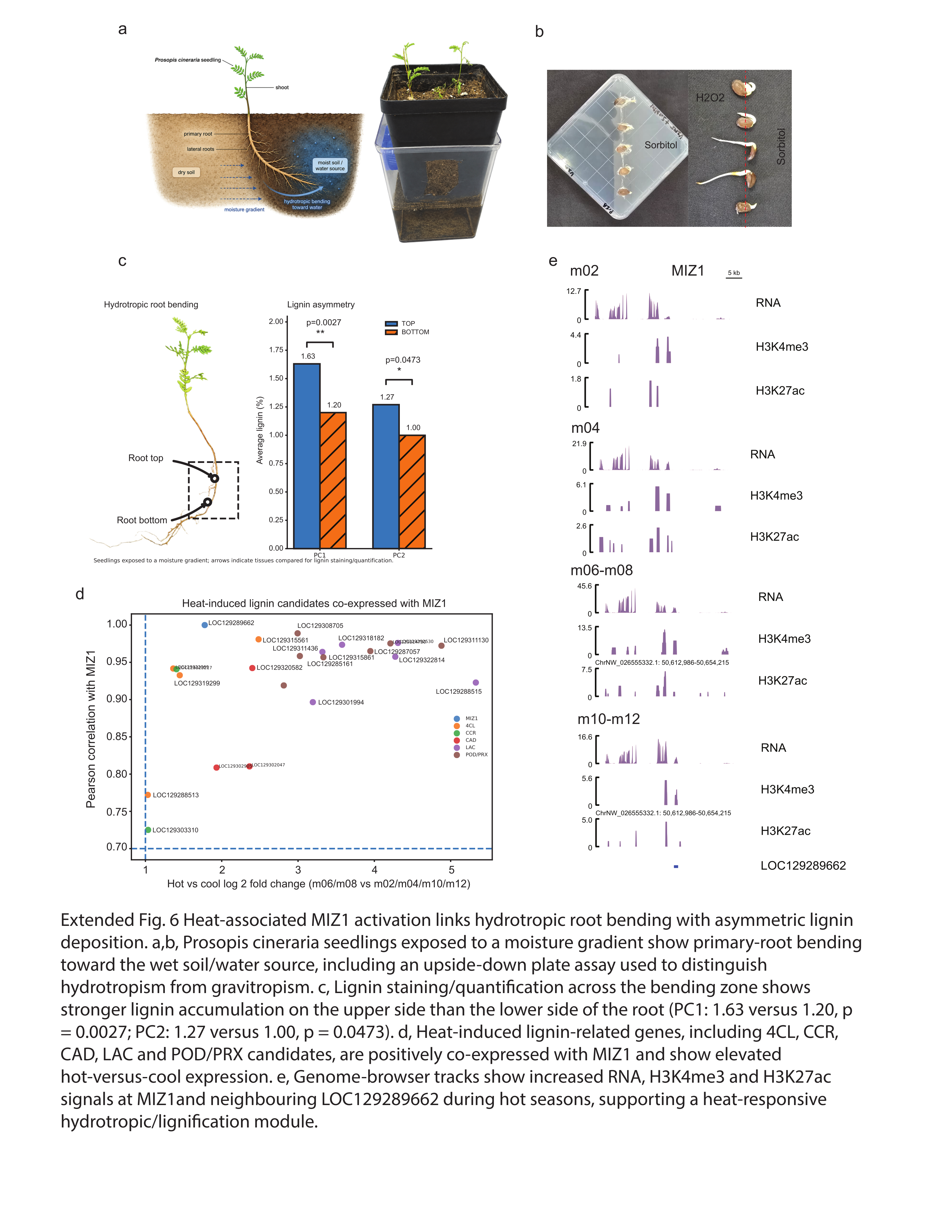

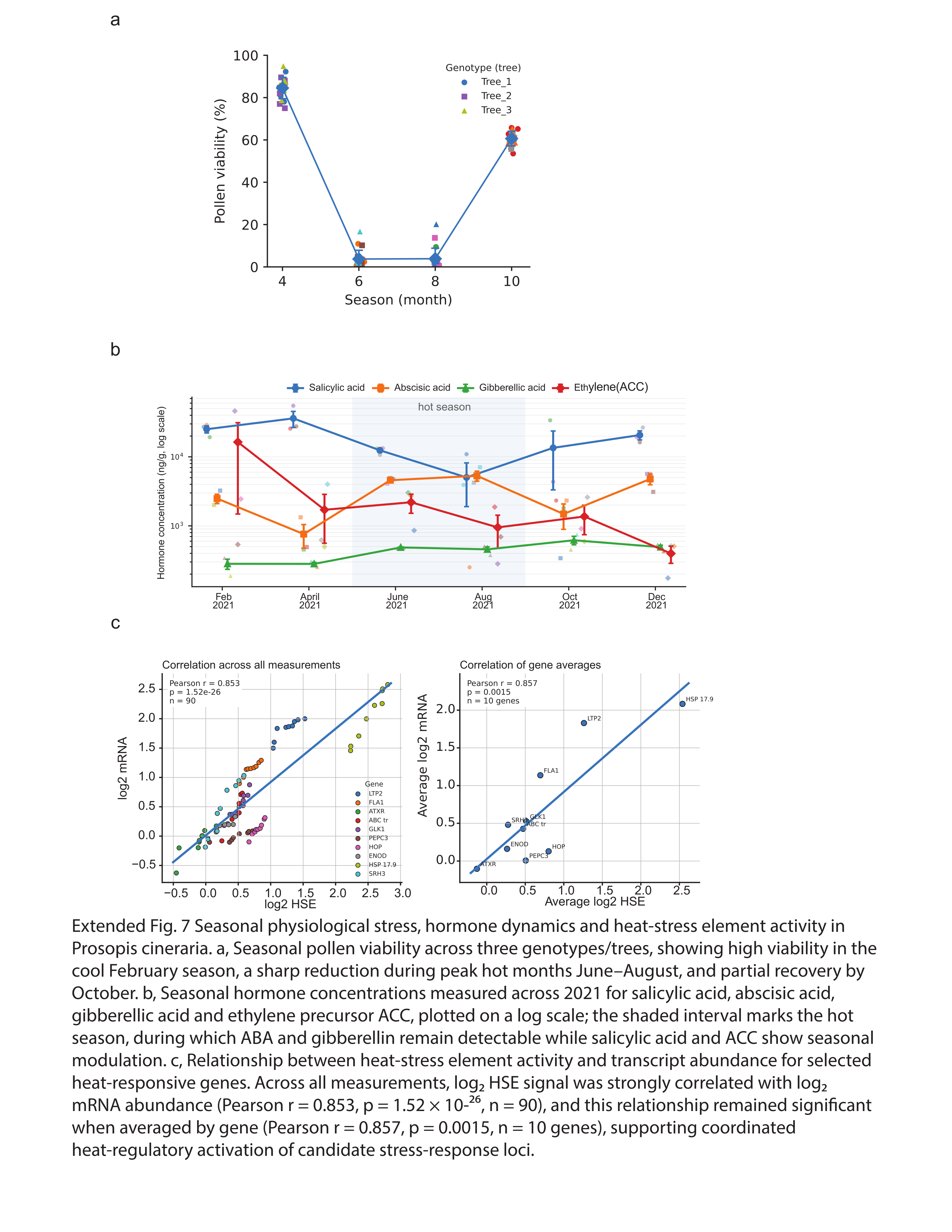

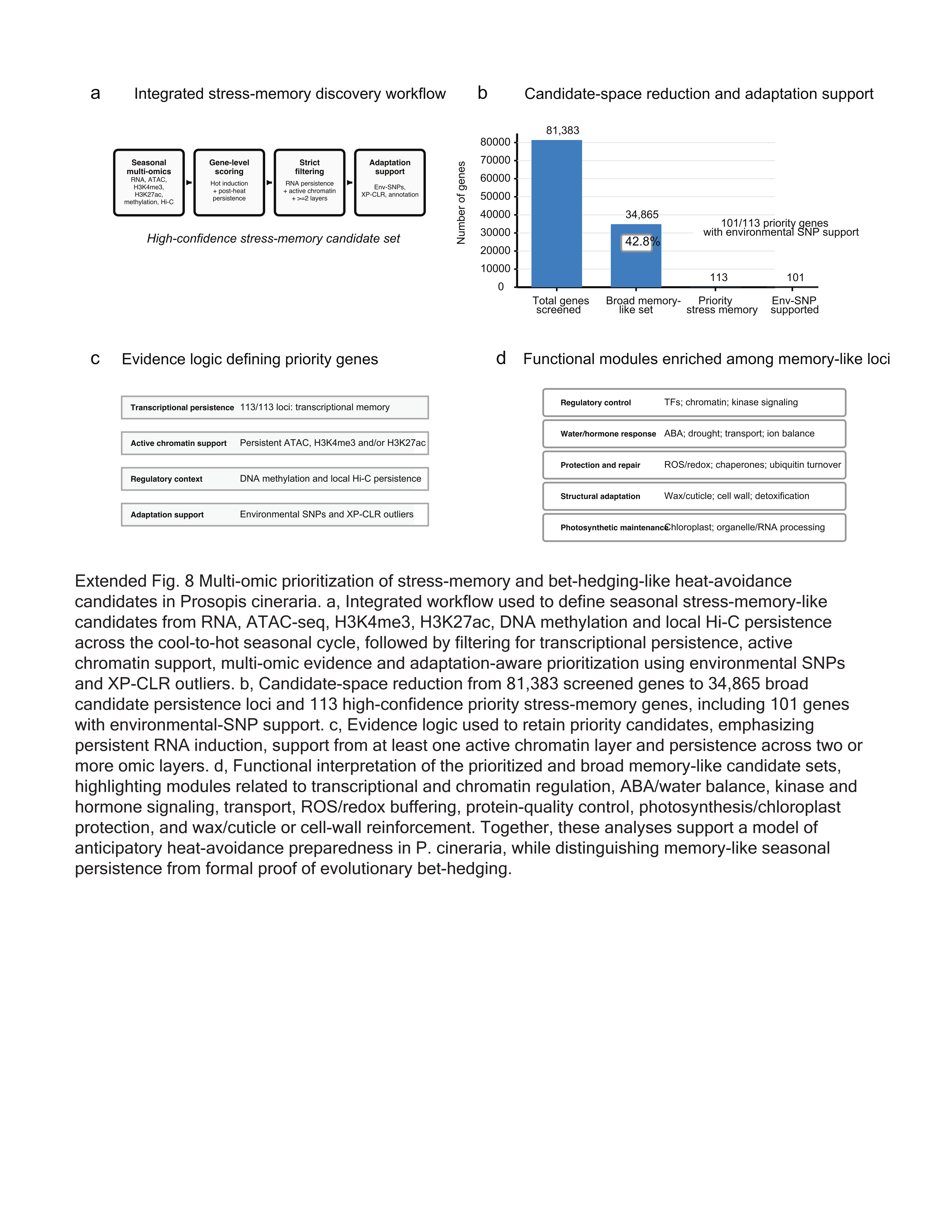

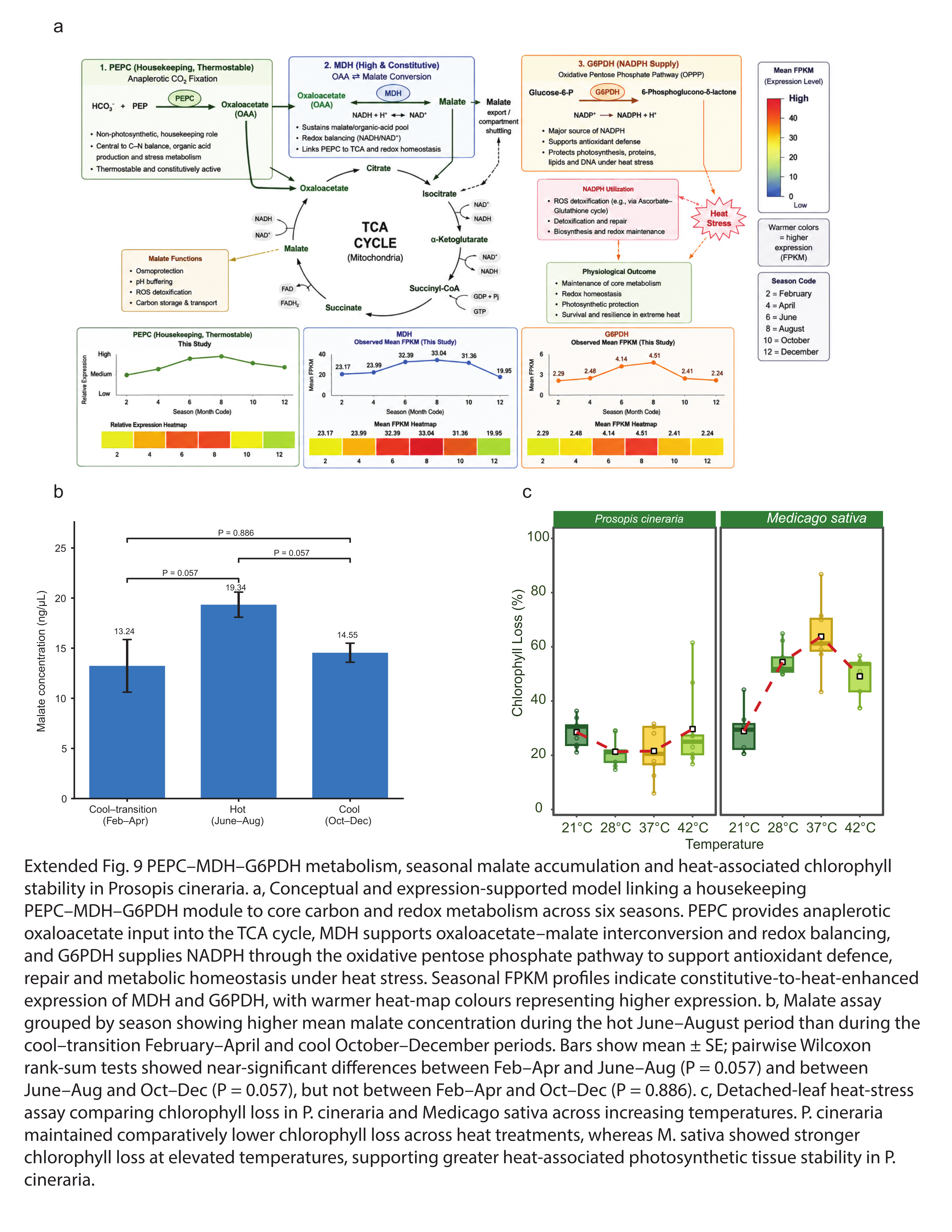

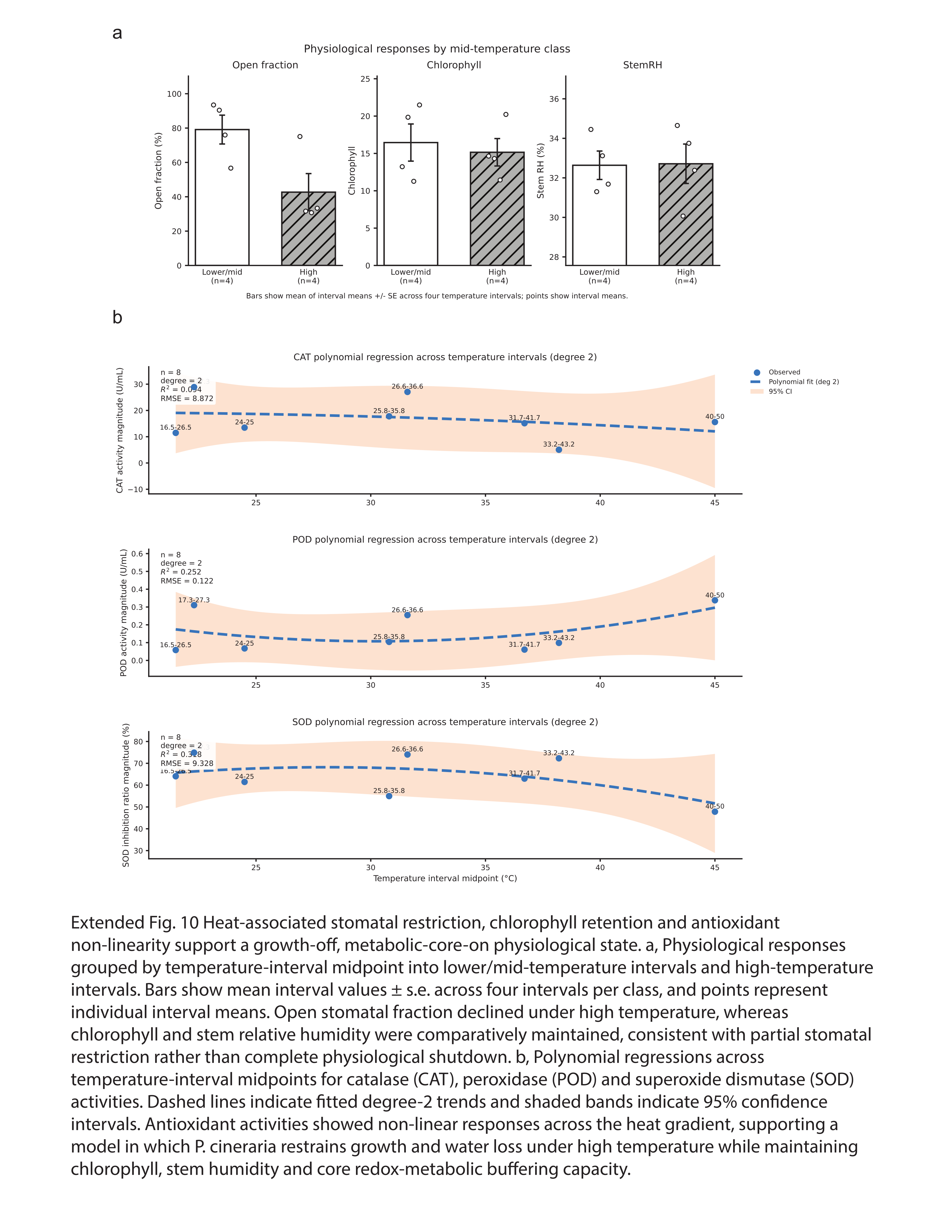
