## Supplementary material for "Reversible chromatin remodeling enables *Prosopis cineraria* survival under recurrent heat extremes": Materials and Methods

**nature climate change**

**Methods**

**Table of Contents**

A. Methods	2

M1. Study Design and Sample Collection

M2. Growth-Chamber Validation Experiment

M3. Multi-Omics Experimental Protocols

M3.1 Hi-C Library Construction

M3.2 Whole-Genome Bisulfite Sequencing (WGBS)

M3.3 ATAC-seq

M3.4 ChIP-seq

M3.5 Transcriptome Sequencing

M4. Computational Data Processing

M4.1 Reference Genome and Annotation

M4.2 Hi-C Data Processing Pipeline

M4.3 WGBS Data Processing and Methylation Analysis

M4.4 ATAC-seq Data Processing

M4.5 ChIP-seq Data Processing

M4.6 Transcriptome Processing and lncRNA Identification

M4.7 Time-Series Clustering of RNA-seq and ATAC-seq Data

M4.8 Multi-Assay Promoter Correlation Analysis

M4.9 Promoter-State Classification and Repeat Annotation

M5. Population and Landscape Genomics

M5.1 Whole-Genome Resequencing and Variant Calling

M5.2 Population Structure and Genetic Differentiation

M5.3 Landscape Genomics

M6. Integrated Multi-Omics Analyses

M6.1 Stress-Memory Candidate Identification and Prioritization

M6.2 Regulatory Network Visualization and Cluster Mapping

M6.3 Post-FIMO Regulatory-Loop Integration and Network Export

M6.4 Machine-Learning Benchmarking for Seasonal Classification

M7. Phenotypic and Biochemical Assays

M7.1 Phenotypic and Physiological Measurements

M7.2 Antioxidant Enzyme Assays

M7.3 Lignin Quantification

M7.4 Epicuticular Wax Quantification

M7.5 Quantitative RT-PCR Validation

M7.6 Root HydrotropiM Assays and Lignin Asymmetry Analysis

M7.7 Pollen Viability, Phytohormone Profiling, and F2 Validation

M7.8 Metabolite Quantification and Comparative Leaf Heat-Stress Assay

References

**A. Methods**

### M1. Study Design and Sample Collection

Leaf material of Prosopis cineraria was collected from trees aged more than 20 years growing in the desert of the Sweihan area (24°17'18.3"N, 55°43'36.2"E), Al Ain, United Arab Emirates. Sampling was conducted in triplicate at alternate-month intervals spanning a full calendar year (January–December 2021), yielding 18 samples in total. Leaf tissue was recovered directly from mature trees, immediately flash-frozen in liquid nitrogen, and transported on dry ice to preserve RNA and chromatin integrity. Each sample was subjected to four complementary sequencing strategies: poly(A)-capture transcriptome sequencing to characterize both coding and non-coding RNA expression across seasons; assay for transposase-accessible chromatin with sequencing (ATAC-seq) to map DNA accessibility for transcriptional regulation; chromatin immunoprecipitation sequencing (ChIP-seq) targeting the histone modifications H3K4me3 and H3K27ac; and whole-genome bisulfite sequencing (WGBS) for genome-wide cytosine methylation profiling (Table M1). To assess the reproducibility of seasonal molecular patterns identified in 2021, samples were collected from the same three trees at six time points during 2025 and subjected to poly(A)-capture transcriptome sequencing.

**Table M1.** Sample description for the multi-omics study of Prosopis cineraria.

| **Sampling Period** | **Transcriptome Sequencing** | **ATAC-seq** | **ChIP-seq (H3K4me3)** | **ChIP-seq (H3K27ac)** | **WGBS** |
| --- | --- | --- | --- | --- | --- |
| February 2021 | 3 samples from 3 different trees. Each sample has two biological replicates and one technical replicate. | 3 samples from 3 different trees | 3 samples from 3 different trees | 3 samples from 3 different trees | 3 samples from 3 different trees |
| April 2021 | 3 samples from 3 different trees. Each sample has two biological replicates and one technical replicate. | 3 samples from 3 different trees | 3 samples from 3 different trees | 3 samples from 3 different trees | 3 samples from 3 different trees |
| June 2021 | 3 samples from 3 different trees. Each sample has two biological replicates and one technical replicate. | 3 samples from 3 different trees | 3 samples from 3 different trees | 3 samples from 3 different trees | 3 samples from 3 different trees |
| August 2021 | 3 samples from 3 different trees. Each sample has two biological replicates and one technical replicate. | 3 samples from 3 different trees | 3 samples from 3 different trees | 3 samples from 3 different trees | 3 samples from 3 different trees |
| October 2021 | 3 samples from 3 different trees. Each sample has two biological replicates and one technical replicate. | 3 samples from 3 different trees | 3 samples from 3 different trees | 3 samples from 3 different trees | 3 samples from 3 different trees |
| December 2021 | 3 samples from 3 different trees. Each sample has two biological replicates and one technical replicate. | 3 samples from 3 different trees | 3 samples from 3 different trees | 3 samples from 3 different trees | 3 samples from 3 different trees |

### M2. Growth-Chamber Validation Experiment

A controlled growth-chamber experiment was conducted to reproduce the temperature conditions recorded during the 2021 field-sampling year. Plants were maintained in a PHcbi MLR-352H chamber at approximately 60% relative humidity with constant weekly irrigation (200 ml, delivered in two applications per week). After an initial acclimatization phase at 25/25 °C (day/night) for three days, plants were sequentially exposed to the temperature and photoperiod regimes corresponding to each sampled month (February, April, June, August, October, and December), as defined by the 2021 environmental profile (Fig. 1). At the completion of the sequential seasonal simulation, plants were subjected to a terminal heat-shock treatment (40–50 °C for three days) to evaluate the induction of heat-responsive transcripts and to investigate potential stress-memory effects. Leaf and root samples were collected at the end of each simulated monthly regime for downstream molecular and phenotypic analyses.

### M3. Multi-Omics Experimental Protocols

#### M3.1 Hi-C Library Construction

Hi-C library preparation and high-throughput sequencing were carried out by Novogene Science and Technology Co., Ltd (Beijing, China). Fresh plant tissues were first ground under liquid nitrogen, then fixed with 1% formaldehyde in vacuum at room temperature for 0.5–1 h. Formaldehyde was quenched by adding 0.15 M glycine under vacuum for 5 min, after which fixed nuclei were collected by filtering through a 40-nm cell strainer. Chromatin was extracted and digested for 16 h with 200 U DpnII restriction enzyme (Qiagen, catalogue no. 59971) at 37 °C. DpnII and MboI share the GATC recognition sequence and are both commonly used in in situ Hi-C protocols; the workflow was designed to allow either enzyme to be substituted depending on library requirements (Belaghzal et al., 2017). After digestion, DNA ends were labelled with biotin and incubated at 37 °C for 45 min, followed by heat inactivation of the enzyme. T4 DNA ligase was added for proximity ligation (1–2 h), after which proteinase K was added to reverse crosslinks (65 °C, 1–2 h). Purified DNA fragments were dissolved in 86 µl of water, unligated ends were removed, and fragments were sized to 350–500 bp. Biotin-labelled ligation products were captured on Dynabeads M-280 Streptavidin (Life Technologies, catalogue no. 60210). Libraries were quality-checked and sequenced on an Illumina NovaSeq 6000 platform with 150 bp paired-end reads (Lieberman-Aiden et al., 2009). Raw reads were pre-filtered to remove: (i) reads containing more than five unidentified nucleotides; (ii) read pairs aligning to adapters; and (iii) reads in which more than 50% of bases had a Phred quality score below 5.

#### M3.2 Whole-Genome Bisulfite Sequencing (WGBS)

DNA extraction, bisulfite conversion, library preparation, and sequencing were performed by Franklin (Hebei Xiong'an) Biotech Co., Ltd (Xiong'an, China). Total genomic DNA was isolated from each sample using the Hi-DNAsecure Plant Kit (Tiangen Biotech, Beijing; catalogue no. DP350) according to the manufacturer's instructions, yielding a minimum of 2,000 µg per sample. DNA integrity and size were verified by electrophoresis on a 0.8% agarose gel. Purified DNA was sonicated (JY92-IIN, Xinzhi, Ningbo) to a fragment-size range of 200–500 bp, and 1 µg of fragmented DNA was subjected to bisulfite conversion using the EZ DNA Methylation-Gold Kit (Zymo Research; catalogue no. D5005). Sequencing libraries were prepared with the Accel-NGS Methyl-Seq DNA Library Kit (Swift Biosciences; catalogue no. 30024), and PCR products of 300–500 bp were enriched, quantified, and sequenced on an Illumina NovaSeq 6000 instrument.

#### M3.3 ATAC-seq

ATAC-seq experiments and sequencing were performed by Franklin (Hebei Xiong'an) Biotech Co., Ltd (Xiong'an, China). Briefly, 500 mg of frozen plant tissue was ground in liquid nitrogen using a tissue lyser. The resulting powder was resuspended in cell lysis buffer, and nuclei were isolated by density-gradient centrifugation for 30 min at 120,000 × g in 60% Percoll/2.5 M sucrose solution. Transposition and library preparation for Illumina sequencing were performed using the TruePrep DNA Library Prep Kit V2 (Vazyme; catalogue no. TD501). Enriched libraries were quantified and sequenced on an Illumina NovaSeq 6000 in 150-bp paired-end mode.

#### M3.4 ChIP-seq

One hundred milligrams of leaf tissue per sample was washed twice in cold PBS, then crosslinked with 1% formaldehyde for 10 min at room temperature. Crosslinking was quenched by adding glycine to a final concentration of 125 mmol/l. Cells were lysed in 50 mM Tris-HCl (pH 8.0) supplemented with 10 mM EDTA, 1% SDS, and 1× protease inhibitor cocktail, and chromatin was isolated on ice. Chromatin was sonicated to yield soluble DNA fragments averaging 200–500 bp. A 20-µl aliquot was retained at −20 °C as input DNA, and 100 µg of chromatin was used per immunoprecipitation. Antibodies (10 µg each) specific to H3K4me3 or H3K27ac were incubated with chromatin overnight at 4 °C, after which 30 µl of protein beads were added for a further 3 h. Beads were washed sequentially with low-salt buffer (20 mM Tris-HCl pH 8.1, 50 mM NaCl, 2 mM EDTA, 1% Triton X-100, 0.1% SDS), high-salt LiCl buffer (10 mM Tris-HCl pH 8.1, 250 mM LiCl, 1 mM EDTA, 1% NP-40, 1% deoxycholic acid), and twice with 1× TE buffer. Immunoprecipitated material was eluted in 300 µl elution buffer (100 mM NaHCO3, 1% SDS), treated with RNase A (8 µg/ml, 6 h at 65 °C) and then proteinase K (345 µg/ml, overnight at 45 °C). Sequencing libraries were prepared using the NEXTFLEX ChIP-Seq Library Prep Kit (Bio Scientific; catalogue no. NOVA-5143-02) and sequenced on an Illumina NovaSeq 6000 in 150-bp paired-end mode.

#### M3.5 Transcriptome Sequencing

Total RNA was extracted from 54 Prosopis samples using the TRIzol method. Strand-specific sequencing libraries were constructed after ribosomal RNA (rRNA) depletion, a strategy chosen to capture both polyadenylated and non-polyadenylated long non-coding RNAs (lncRNAs), as some lncRNAs harbour poly(A) tails while others do not. rRNA-depleted RNA was fragmented randomly; first-strand cDNA was synthesized using random hexamer primers, and second-strand cDNA synthesis was performed in the presence of buffer, dNTPs, RNase H, and DNA polymerase I. The cDNA product was purified with a QIAquick PCR purification kit, end-repaired, A-tailed, and ligated to sequencing adapters. UNG enzyme digestion of the second strand preserved strand specificity. Libraries were size-selected by agarose gel electrophoresis, PCR-amplified, and sequenced on an Illumina instrument in paired-end mode to generate transcriptome reads.

### M4. Computational Data Processing

#### M4.1 Reference Genome and Annotation

All analyses used the Prosopis cineraria reference genome assembly GCF_029017545.1 (AM2901754v1), generated in-house and deposited in NCBI in 2022. A FASTA index was built with SAMtools faidx, and chromosome sizes were extracted from the resulting .fai file. Genome annotations were derived from the corresponding GFF file; gene features were converted from 1-based closed GFF coordinates to 0-based half-open BED coordinates, and transcription start sites (TSSs) were inferred from strand orientation (start coordinate for positive-strand genes; end coordinate for negative-strand genes). Gene names were taken from the Name attribute of the GFF, with gene and ID attributes used as fallbacks when Name was absent (Li et al., 2009; Danecek et al., 2021).

#### M4.2 Hi-C Data Processing Pipeline

##### M4.2.1 Overview and Computational Environment

Hi-C sequencing data were processed using a reproducible, stage-aware command-line workflow (run_hic_pipeline.sh) that converts paired-end FASTQ files into per-season chromatin contact maps in both multi-resolution cooler (.mcool) and Juicebox-compatible (.hic) formats. The pipeline was executed on a high-performance computing cluster using module-managed software: FastQC v0.11.9, fastp v0.23.2, bwa-mem2 v2.3, SAMtools v1.18, MultiQC v1.33, pairtools v1.1.3, cooler v0.10.4, and Singularity v4.2.1. Sample metadata were provided in a tab-delimited samples.tsv file. The workflow maintained a structured directory tree for quality-control reports, trimmed reads, alignments, pairs files, cooler matrices, Juicer outputs, logs, reference files, genome tracks, and plots. Staged re-analysis was supported via RERUN_STAGE and force-rerun flags, allowing individual steps to be repeated without overwriting completed outputs.

##### M4.2.2 Read Quality Control and Alignment

Raw reads were first assessed with FastQC to evaluate per-base quality, base composition, sequence duplication, and other common sequencing artefacts (Andrews, 2010). When trimming was enabled, paired reads were processed with fastp using paired-end adapter detection, tail-quality trimming at a mean threshold of Q20, a minimum retained read length of 30 bp, and eight processing threads; both HTML and JSON reports were retained for auditability (Chen et al., 2018). Trimmed reads were aligned to the indexed reference genome using bwa-mem2 with 16 threads, an accelerated implementation of the BWA-MEM algorithm that maintains full compatibility with downstream SAM/BAM tools (Vasimuddin et al., 2019). Alignments were converted to coordinate-sorted, indexed BAM files using SAMtools (Li et al., 2009; Danecek et al., 2021). A secondary query-name-sorted BAM was produced for each sample to facilitate paired-read grouping required by pairtools.

##### M4.2.3 Contact Pair Extraction, Sorting, and Deduplication

Hi-C contact pairs were extracted from query-name-sorted BAM files using pairtools parse, specifying the reference chrom.sizes file, a minimum mapping quality of 20, the 5-unique walks policy, and exclusion of SAM records from the output stream. Parsed contacts were sorted with pairtools sort and deduplicated with pairtools dedup to remove PCR and optical duplicate ligation products (Open2C et al., 2024b). Deduplicated contacts were written as compressed .pairs.gz files; duplicate pairs were retained separately as .dups.pairs.gz files. The resulting .pairs files are compatible with Pairix indexing and downstream matrix construction (Lee et al., 2022). For each season, pairtools stats was used to summarize cis and trans contact counts. Distance-decay and cis/trans contact plots were generated using a custom Python script; when pairtools distance-bin statistics were unavailable, up to 5,000,000 contact pairs were sampled from the .pairs file to compute genomic distance distributions directly. Excessive trans contacts, weak distance decay, or abnormal duplicate rates are recognized indicators of library or processing failure and were used as quality-exclusion criteria (Lieberman-Aiden et al., 2009; Belaghzal et al., 2017). Per-sample FastQC reports, pairtools statistics, and QC plots were aggregated with MultiQC (Ewels et al., 2016).

##### M4.2.4 Matrix Generation and Multi-Resolution Balancing

Deduplicated .pairs.gz files were converted into binned contact matrices using cooler at a base resolution of 5 kb (Abdennur and Mirny, 2020). Where available, pairix was used to index the compressed pairs file before matrix loading (cooler cload pairix); if indexing failed, the workflow fell back to cooler cload pairs with explicit column definitions. Each 5-kb base matrix was then multi-resolution balanced with cooler zoomify, generating .mcool files at 5, 10, 25, 50, 100, 250, and 500 kb and 1 Mb resolutions. Iterative correction/balancing normalizes for systematic coverage biases arising from restriction-fragment length, GC content, and mappability (Imakaev et al., 2012). In parallel, Juicebox-compatible .hic files were generated for each season by processing the same FASTQ inputs through Juicer inside a Singularity container, using the Prosopis reference FASTA, chromosome sizes, restriction-site file, and 16 processing threads (Durand et al., 2016).

##### M4.2.5 Matrix-Level Quality Control of Seasonal Hi-C Contact Maps

Seasonal Hi-C libraries were stored as multi-resolution cooler files (.mcool) and interrogated at 50-kb resolution using a custom Python workflow operating on sparse contact matrices. For each season, we quantified total non-zero pixels, total contact counts, the fraction of bins retaining balancing weights, and the fraction of bins classified as bad or unbalanced (Abdennur and Mirny, 2020; Imakaev et al., 2012). Contacts across the top 14 scaffolds were partitioned into cis (intra-chromosomal) and trans (inter-chromosomal) classes, and cis contacts were further subdivided into short-, mid-, and long-range bins. Contact probability P(s) decay curves were calculated by aggregating contact frequency over increasing genomic distances (Lieberman-Aiden et al., 2009; Rao et al., 2014; Yardimci et al., 2019). Quality summaries were visualized as bar plots of cis/trans fractions, genomic distance-decay curves, and representative chromosome-scale heat maps at 50 kb for scaffold NW_026555332.1. Libraries were considered acceptable when they exhibited strong diagonal enrichment, Mooth monotonic P(s) decay, predominance of cis over trans contacts, and broadly consistent bin balanceability (Lieberman-Aiden et al., 2009; Rao et al., 2014; Yardimci et al., 2019). Detailed results from this quality assesMent are presented in Results SR1.

##### M4.2.6 Cis-Loop and Interchromosomal Contact Analysis at 10 kb

Seasonal Hi-C contact maps were analysed at 10-kb resolution for fine-scale loop calling and interchromosomal screening. For each season, the 10-kb matrix was extracted from the multi-resolution .mcool file using the cooler Python API (Abdennur and Mirny, 2020). If a bin-level weight column was absent, matrix balancing was performed with cooler balance (12 threads). Distance-dependent cis background was estimated using cooltools expected-cis with 12 threads and sample-specific chromosome view files (Open2C et al., 2024a). Cis-enriched focal contacts were identified with cooltools dots using a maximum cis separation of 2 Mb, a tile size of 6 Mb, a clustering radius of 40 kb, and an FDR threshold of 0.02 (Rao et al., 2014; Open2C et al., 2024a). Dot calls were exported in BEDPE format for each season. Enriched interchromosomal candidate contacts were identified from balanced 10-kb matrices by computing, for each trans pixel, the balanced contact signal (raw count × weight1 × weight2), and calculating fold enrichment over the chromosome-pair median as well as conventional and robust z-scores. Trans candidates were retained when fold enrichment exceeded 5× the chromosome-pair median and the conventional z-score was ≥3.0; the top 500 candidates per chromosome pair were exported. Because this screen uses a custom enrichment filter rather than a formally calibrated loop caller, trans contacts were interpreted as candidates requiring downstream biological validation across seasons (Rao et al., 2014; Durand et al., 2016).

##### M4.2.7 Seasonal TAD Boundary Inference and Temporal Comparison

TAD boundaries were inferred independently for each season from balanced .mcool matrices using cooltools insulation with a 500-kb diamond window, Li thresholding for boundary selection, and the balanced matrix weight column (Crane et al., 2015; Open2C et al., 2024a). The resulting insulation tables were post-processed to extract cleaned boundary BED files and reconstructed TAD intervals. Adjacent boundary calls within one bin were merged, retaining the strongest local signal. TADs were reconstructed as intervals between consecutive boundary anchors on each chromosome; domains shorter than three bins were merged into adjacent domains, and unusually large domains were annotated relative to a soft upper bound of 200 bins. Temporal TAD dynamics were then analysed with the TimeCompare function of the TADCompare R package at 50-kb resolution, using sparse matrices exported from the .mcool files (Cresswell and Dozmorov, 2020). Boundary changes classified by TADCompare include shifts, gains, losses, merges, and splits. Regions with fewer than two available seasonal matrices were skipped. A combined table of TimeCompare-derived boundaries and a summary recording region, number of seasons used, and consensus score were produced for each chromosome.

##### M4.2.8 A/B Compartment Calling and Differential Analysis

A/B compartments were called for each season using cooltools eigs-cis, requesting the first three cis eigenvectors from balanced .mcool matrices at 50-kb resolution (Lieberman-Aiden et al., 2009; Imakaev et al., 2012; Open2C et al., 2024a). Eigenvector sign was anchored by a gene-density phasing track so that positive eigenvector-1 values corresponded to gene-rich, A-like chromatin. Bins with eigenvector-1 ≥0 were assigned compartment A; negative values were assigned compartment B. Differential compartment analysis was performed with dcHiC using matrices exported from the .mcool files (Chakraborty et al., 2022). A dcHiC input table paired each seasonal matrix with its corresponding bin BED file. The wrapper executed dcHiC in three steps: cis principal-component analysis, PC selection, and differential compartment analysis, using quantile-normalized compartment scores and multivariate distances to identify significant compartmentalization changes across seasons. A dedicated summary script combined insulation boundaries, reconstructed TADs, and A/B compartment calls across seasons, computing boundary recurrence with a 50-kb positional tolerance and classifying TAD transitions (stable, expanded, contracted, gained, lost, merged, split) using reciprocal-overlap criteria.

To quantify compartment-activity coupling (Extended Data Fig. 3), median assay signal in A and B compartments was compared genome-wide and within promoter/TSS ±2-kb intervals for each season. Spearman correlations were calculated between eigenvector-1 scores and RNA, H3K4me3, and H3K27ac signal values per bin, providing a continuous measure of the compartment-transcription relationship across the seasonal series. A-minus-B signal differences were computed for each assay at both 50-kb bin and promoter-window resolution to track seasonal changes in the compartment-activity contrast. This approach follows established Hi-C compartment frameworks and plant 3D-genome applications (Lieberman-Aiden et al., 2009; Dixon et al., 2012; Ouyang et al., 2020; Tourdot et al., 2023).

##### M4.2.9 H3K4me3 Integration with Compartment Dynamics

H3K4me3 signal was integrated with seasonal compartment dynamics at three genomic scales: 50-kb compartment bins, annotated gene bodies, and promoter intervals. H3K4me3 bedGraph files were matched to seasonal labels (m02, m04, m06, m08, m10, m12) and signal was aggregated as overlap-weighted mean values across bins, gene bodies, and promoters (defined as 2 kb upstream to 500 bp downstream of the TSS; orientation reversed for negative-strand genes). Replicate profiles were averaged within each season. For each adjacent seasonal transition, the workflow computed signal in season 1, signal in season 2, and the delta value (season 2 − season 1). Pairwise class comparisons used permutation tests on delta H3K4me3 values (5,000 permutations; minimum class size of 20; plus-one P-value correction). An auxiliary script restricted the same framework to the top-14 scaffolds and identified B-to-A promoter-gain genes (any positive gain, top 25%, or top 10%) and A-to-B genes (all, or filtered by bottom 25% or 10% promoter loss). Where an InterPro annotation file was provided, functional annotations were merged using gene or LOC identifiers (Jones et al., 2014).

#### M4.3 WGBS Data Processing and Methylation Analysis

Genome-wide cytosine methylation was analysed across 18 WGBS libraries, with an average yield of 24.26 GB per sample and a mean mapping rate of 86.51%. Raw reads were generated by CASAVA base calling and quality-filtered by removing adapter-contaminated reads, reads with more than 5% ambiguous bases, and reads in which more than 20% of bases had quality scores below 10. Clean paired-end reads were aligned to the Prosopis reference genome using BMAP v2.90 with parameters -m 30 -x 700 -u -v 8 -z 33 -p 4 -n 0 -w 20 -s 16 -f 10 -L 150 (Xi and Li, 2009). Alignments were coordinate-sorted and indexed with SAMtools (Li et al., 2009). Methylation levels at individual cytosines were defined as Nm/(Nm + Nnm), where Nm is the count of reads supporting a methylated cytosine and Nnm is the count supporting an unmethylated cytosine. Downstream methylome characterization encompassed coverage statistics for all cytosine contexts (CG, CHG, CHH); methylation levels across genomic elements (mRNA, exon, intron, ncRNA); methylation profiles across seven transcriptional element classes (upstream, downstream, first and last exon, first intron, and other exon and intron); chromosome-level summaries; the relative contributions of mCG, mCHG, and mCHH to total methylated cytosines; global methylation level distributions; sequence-bias entropy analysis across ±9 nucleotide windows surrounding methylated sites; heat maps of methylation relative to genomic regions and CpG density; and binned mean methylation profiles across gene-associated elements with the TSS marked explicitly. Differentially methylated regions (DMRs) were called using the R package DSS v2.40.0 with thresholds of delta ≥0.1 and adjusted P ≤0.05 (Park and Wu, 2016).

#### M4.4 ATAC-seq Data Processing

Chromatin accessibility was profiled across 18 ATAC-seq libraries. Raw reads were quality-filtered with fastp: adapter sequences were trimmed, reads containing undetermined bases were removed, and reads with Q ≤5 in ≥50% of positions were excluded (Chen et al., 2018). The resulting high-quality reads (Q20 ≈98%; Q30 ≈94–95%) were aligned to the reference genome using Bowtie2 (Langmead and Salzberg, 2012), and BAM files were converted to bigWig tracks for visualization in IGV or the UCSC Genome Browser. Alignment-level quality control was performed with deepTools (Ramirez et al., 2016), encompassing insert fragment-length distributions, gene-coverage heat maps with gene bodies scaled to a uniform length and 3-kb flanking regions displayed, sequencing-depth statistics based on randomly selected 1-Mb windows, PCA on genome-wide coverage profiles, and pairwise Pearson correlation analysis among samples. Accessible chromatin peaks were called with MACS2 (Zhang et al., 2008) and annotated with the R package ChIPseeker, which assigned each peak to promoters, UTRs, introns, and other genomic features and recorded nearest genes and TSS distances. For consensus peak calling, BAM files from triplicates within each time point were merged using SAMtools v1.18 (Li et al., 2009; Danecek et al., 2021), and MACS2 v2.2.9.1 was run with --call-summits --shift -100 --extsize 200 --nomodel -B --SPMR using the reported Prosopis cineraria genome size (691,386,579 bp). Transcription factor footprint scores were calculated from ATAC-seq cut-site signal using TOBIAS v0.17.0 FootprintScores; per-base signal was provided via --signal and MACS2-derived narrowPeak regions via --regions (Bentsen et al., 2020). TOBIAS estimates localized depletion of Tn5 insertion signal within accessible regions relative to flanking signal, consistent with the protection pattern generated by DNA-bound regulatory proteins; these signals were therefore interpreted as candidate regulatory footprints rather than direct evidence of TF binding. For motif enrichment, peak lists were sorted by fold enrichment, the top 1,000 peak sequences were extracted, and de novo and known motif analyses were performed using HOMER (Heinz et al., 2010). Known-motif enrichment results across samples were consolidated into a unified TF-by-sample matrix by converting natural-log P-values to −log10 P-values.

#### M4.5 ChIP-seq Data Processing

ChIP-seq was performed on 144 libraries comprising immunoprecipitated and matched input samples for H3K27ac and H3K4me3. Raw paired-end reads were quality-filtered with fastp to remove adapters, reads with undetermined bases, and low-quality reads (Q ≤5 in >50% of positions) (Chen et al., 2018). Clean reads were aligned to the reference genome using Bowtie2 (Langmead and Salzberg, 2012), and bigWig tracks were generated for visualization. BAM files were processed with SAMtools v1.3.1, including removal of PCR duplicates where applicable (Li et al., 2009). Alignment quality was assessed from mapping statistics (total, unique, multi-mapped, and discordant reads) and from deepTools summaries: insert fragment-length distributions, sequencing-depth assesMent, gene-body coverage profiles across ±3-kb flanking regions, PCA on genome-wide coverage, and Pearson inter-sample correlations (Ramirez et al., 2016). Peaks were called using MACS2 v2.1.1.20160309 against the matched input library for each immunoprecipitated sample (bandwidth = 300 bp; model fold = 5 and 50; q-value cutoff = 0.05) (Zhang et al., 2008). Peaks were assigned to the gene whose TSS was nearest the peak midpoint, and genomic distributions were further annotated with ChIPseeker into promoter, UTR, exon, intron, downstream, and intergenic categories.

#### M4.6 Transcriptome Processing and lncRNA Identification

Transcriptome profiling was performed on 54 strand-specific RNA-seq libraries with an average yield of 13.22 Gb per sample and a mean genome-mapping rate of 97.04%. To minimize ribosomal contamination, clean reads were first aligned to an rRNA database using SOAP2 v2.2.1 (-m 0 -x 1000 -s 40 -l 32 -v 5 -r 2 -p 3), and only rRNA-depleted reads were retained. Filtered reads were mapped to the Prosopis reference genome using HISAT2 v2.2.1 (--phred33 --sensitive --no-discordant --no-mixed -I 1 -X 1000 --rna-strandness RF) (Kim et al., 2019). Transcripts were reconstructed per sample with StringTie v2.2.1 (--rf -f 0.3 -j 3 -c 5 -g 100 -s 10000 -p 8) (Pertea et al., 2015), merged with StringTie-merge v2.2.1 (-m 200 -c 3), and compared against the reference annotation with gffcompare v0.12.6 to identify novel transcripts. Candidate lncRNAs were selected by requiring transcript coverage ≥3, transcript length ≥200 bp, novelty relative to the existing annotation, and non-coding status confirmed by at least two of three tools: CPC v0.9-r2 (Kong et al., 2007), CNCI v2 (-m ve -p 4) (Sun et al., 2013), and Pfam domain scanning with PfamScan v1.6 (-cpu 4 -e_seq 0.001 -e_dom 0.001) (Jones et al., 2014). Expression levels of mRNAs and lncRNAs were quantified using Ballgown v2.26.0 with FPKM values derived from StringTie output (-e -B) (Frazee et al., 2015). Sample similarity was assessed by pairwise Pearson correlation and PCA. Differential expression analysis used four algorithms depending on replicate structure: DESeq2 v1.34.0 (fold change ≥2, adjusted P ≤0.05) (Love et al., 2014), EBSeq v1.34.0 (fold change ≥2, PPEE ≤0.05), NOISeq v2.14.1 (fold change ≥2, probability ≥0.8), and PoissonDis v1.0 for comparisons without biological replicates (log2FC >1, FDR <0.001). Differentially expressed genes were further characterized by hierarchical clustering, K-means clustering over multiple time points, GO enrichment (topGO v2.46.0; corrected P <0.05), KEGG pathway analysis, protein–protein interaction analysis using the STRING database and Cytoscape (Shannon et al., 2003), and transcription factor prediction using PlantTFDB. For lncRNAs, family assignment used Infernal v1.1.4; antisense interactions were predicted with RNAplex v2.5.0; cis targets were defined as protein-coding genes within 100 kb of differentially expressed lncRNAs; and trans targets were identified using cor.test() in R (r >0.95, P <0.05, minimum five samples).

To validate the reproducibility of seasonal expression states, RNA-seq profiles from the 2021 multi-omic reference year were compared with an independent 2025 validation dataset sampled from the same three trees at the same six time points (Extended Data Fig. 2). Expression values from both years were normalized to log2-transformed FPKM values and compared per season using Pearson correlation (linear concordance) and Spearman rho (rank-order consistency). Genes were stratified by expression quartile to determine whether cross-year agreement was restricted to highly expressed transcripts or reflected a broad seasonal programme. This validation follows standard transcriptomic reproducibility practice in which independent time series are used to distinguish recurrent biological seasonality from year-specific variation (Love et al., 2014; Conesa et al., 2016).

#### M4.7 Time-Series Clustering of RNA-seq and ATAC-seq Data

Gene-level FPKM expression values from the RNA-seq analysis were integrated with sample metadata (sample identity, genotype, temperature) and InterProScan-derived functional annotation (Jones et al., 2014). Seasonal identity was encoded as six ordered time points: February (m02), April (m04), June (m06), August (m08), October (m10), and December (m12). Replicate samples from the same season were averaged to generate one seasonal expression profile per gene. Genes were filtered by requiring expression above a minimum abundance threshold in at least a minimum number of seasons; genes with no seasonal variance were excluded. Profiles were standardized prior to clustering so that temporal pattern rather than absolute expression level drove cluster assignment. Soft clustering was then performed using fuzzy c-means as implemented in Mfuzz (Kumar and Futschik, 2007) within a Bioconductor-compatible ExpressionSet framework (Huber et al., 2015). Cluster number was not fixed a priori; instead, candidates were evaluated using average silhouette width, the Calinski-Harabasz index, the Davies-Bouldin index, fuzzy partition coefficient, and fuzzy partition entropy (Calinski and Harabasz, 1974; Davies and Bouldin, 1979; Rousseeuw, 1987). The final number was selected by composite ranking favouring high silhouette width and Calinski-Harabasz values, and low Davies-Bouldin, fuzzy partition entropy values. Heatmaps of standardized cluster patterns were produced with ComplexHeatmap (Gu et al., 2016). Temperature and seasonal associations were tested by Pearson correlation of cluster centroids against numeric temperature and by linear models; P-values were adjusted by Benjamini-Hochberg FDR (Benjamini and Hochberg, 1995). Cluster member genes were annotated against InterProScan-derived LOC-based annotation tables.

Time-course chromatin accessibility dynamics from ATAC-seq, as well as time-course differential expression, were analysed in R using the TCseq package v1.26.0 (Wang et al., 2019). Peak regions were imported from MACS2 narrowPeak files using peakreference to create a unified genomic feature reference. Aligned BAM files were supplied to countReads to quantify read coverage, and differential binding/accessibility was assessed with DBanalysis using February as the baseline against all later time points. Temporal peak patterns were clustered using k-means (k = 4) on RPKM-normalized, standardized values with timeclust, and cluster profiles were visualized with timeclustplot.

#### M4.8 Multi-Assay Promoter Correlation Analysis

To assess the seasonal coupling between transcription and chromatin state (Extended Data Fig. 1), promoter-centred signal matrices were constructed for all expressed genes using TSS ±2-kb intervals. For each assay — RNA-seq, H3K4me3 ChIP-seq, H3K27ac ChIP-seq, ATAC-seq, and CG/CHG/CHH WGBS methylation — per-gene mean signal was summarized over the same TSS-centred windows so that cross-assay comparisons used matched genomic intervals. Pairwise Pearson correlations and Spearman rho were calculated between all assay vectors at each seasonal time point. Selected bivariate relationships were visualized as density/scatter plots with correlation statistics annotated. H3K4me3 and H3K27ac were treated as active promoter and enhancer marks respectively; ATAC-seq was treated as a measure of chromatin accessibility reflecting regulatory competence rather than direct transcriptional output; and DNA methylation contexts were analysed separately because CG, CHG, and CHH methylation have distinct maintenance pathways and regulatory interpretations in plants (Roudier et al., 2011; Creyghton et al., 2010; Buenrostro et al., 2013; Law and Jacobsen, 2010; Zhang et al., 2018).

#### M4.9 Promoter-State Classification and Repeat Annotation

To separate two seasonally distinct repression programmes (Extended Data Fig. 4), promoter states were classified by integrating CHH methylation and H3K4me3 signal across all six seasons. Candidate TE-proximal promoters were annotated by intersecting TSS ±2-kb intervals with repeat/TE annotations from the Prosopis reference; TE classes were summarized to distinguish TIR, LTR, Helitron, and other repeat contributions. High-CHH/low-H3K4me3 promoters were identified as candidate cool-season RdDM-like loci; low-H3K4me3 promoters specifically reduced during m06/m08 and without strong CHH enrichment were designated as the hot-season PRC2-compatible candidate module. Functional labels were assigned from gene annotation and manually grouped into stress/defence, flowering/developmental, and chromatin-regulatory classes. Caution was applied in interpreting these states: CHH methylation maintenance near repeats is commonly mediated by RNA-directed DNA methylation (RdDM), whereas low H3K4me3 during hot seasons is considered PRC2-compatible but not confirmed without direct H3K27me3 profiling (Law and Jacobsen, 2010; Matzke and Mosher, 2014; Mozgova and Hennig, 2015; Xiao and Wagner, 2015; Zhang et al., 2018).

### M5. Population and Landscape Genomics

#### M5.1 Whole-Genome Resequencing and Variant Calling

A whole-genome resequencing dataset comprising 222 Prosopis samples was assembled, combining 13 internally generated samples with publicly available accessions from NCBI BioProject PRJEB82449. After sample-level quality control, 216 individuals were retained for downstream population genomic analyses. Short reads were aligned to the reference genome using BWA-MEM (Li, 2013), followed by coordinate sorting and indexing of BAM files (Li et al., 2009). PCR duplicates were marked with GATK MarkDuplicates, and per-sample variant discovery was performed with GATK HaplotypeCaller in GVCF mode (McKenna et al., 2010). Biallelic SNPs were extracted and filtered, and the resulting biallelic SNP VCF was taken forward for population genomic analyses, following established best practices for short-read variant discovery (Li, 2013; Li et al., 2009; McKenna et al., 2010).

The filtered biallelic SNP VCF was converted to PED/MAP and binary BED/BIM/FAM formats using PLINK (Purcell et al., 2007; Chang et al., 2015). SNPs were filtered at a maximum missingness of 2% (--geno 0.02); individuals with more than 2% missing data were then excluded (--mind 0.02); variants with minor allele frequency (MAF) below 2% were removed (--maf 0.02). Linkage disequilibrium pruning was applied with --indep-pairwise 50 10 0.2 (50-SNP window, 10-SNP step, r2 threshold of 0.2).

#### M5.2 Population Structure and Genetic Differentiation

Population structure was inferred from the LD-pruned dataset using SNPRelate and LEA in R (Zheng et al., 2012; Frichot et al., 2014; Frichot and Francois, 2015). The VCF was converted to GDS format with snpgdsVCF2GDS, and SNPs were filtered within R at MAF ≥0.05 and maximum per-marker missingness ≤0.10 before a second LD-pruning step in SNPRelate (200-kb window; LD threshold r2 ≤0.2). Individual ancestry was estimated using the sNMF algorithm in LEA across K = 1–10 ancestral clusters, with five replicate runs per K, alpha = 100, diploid coding, and multithreaded execution (Frichot et al., 2014). The optimal K was determined from an Evanno-like Delta K summary of cross-entropy values across replicates (Evanno et al., 2005). Each individual was assigned to a hard cluster based on the maximum ancestry coefficient in the best-supported Q-matrix. PCA was performed in SNPRelate on the LD-pruned SNP set; individuals were plotted in PC1/PC2 space coloured by sNMF-derived cluster assignment, with 95% confidence ellipses drawn around inferred groups (Zheng et al., 2012). Pairwise genetic differentiation among inferred clusters was estimated using Weir-Cockerham FST computed with adegenet and hierfstat from allele-coded genotype data (Jombart, 2008; Jombart and Ahmed, 2011; Goudet, 2005; Weir and Cockerham, 1984). Population structure results were summarized as a multi-panel figure combining Delta K, admixture bar plots, a PCA scatter plot, and a pairwise FST heat map.

Cross-population composite likelihood ratio (XP-CLR) analysis was performed as a downstream selection scan on the most highly differentiated cluster pairs identified from FST results (Chen et al., 2010). XP-CLR integrates multilocus allele-frequency differentiation across sliding windows, improving sensitivity for selective sweep detection relative to single-marker comparisons. Candidate regions with elevated XP-CLR scores were interpreted as putative targets of divergent selection between the focal populations.

#### M5.3 Landscape Genomics

Landscape genomics was performed using a filtered VCF of high-quality biallelic SNPs together with an environmental dataset comprising seven standardized predictors: bio03 (isothermality), bio08 (mean temperature of the wettest quarter), bio09 (mean temperature of the driest quarter), bio11 (mean temperature of the coldest quarter), bio13 (precipitation of the wettest month), bio15 (precipitation seasonality), and soil nitrogen. Environmental variables were z-transformed and missing values were imputed. The genotype matrix was converted from VCF using bcftools and custom R scripts.

Redundancy analysis (RDA) was implemented with the vegan R package. A partial RDA model (genotypes ~ environment | MEM) was used to control for spatial autocorrelation, where MEM represents Moran's Eigenvector Maps derived from geographic coordinates. Model significance was evaluated by 999 permutations. SNP loadings from significant RDA axes were extracted, and outliers beyond ±3 SD of the mean loading were designated as candidate loci under selection. Gradient Forest analysis was performed with the gradientForest package using 500 regression trees per SNP (Ellis et al., 2012); environmental variable importance was computed as cumulative split importance, and SNPs in the top 1% importance percentile were classified as GF outliers. Latent factor mixed modelling (LFMM2) was applied to identify SNP–environment associations while accounting for population structure, using the lfmm2 R package with K = 1–7 latent factors (Frichot et al., 2014). The genomic inflation factor λ was computed for each environmental variable; K = 5 minimized λ and was selected for all final LFMM2 runs. P-values were adjusted by genomic control, and Bonferroni-corrected significance thresholds were applied based on total SNP number. SNPs detected as outliers by at least two of three methods (RDA, GF, LFMM2) were classified as high-confidence adaptive candidates and compiled in a combined table. Overlaps were visualized using the VennDiagram package, and Manhattan plots were generated with ggplot2 highlighting multi-method outliers. All analyses were conducted in R v4.3.0 (R Core Team, 2024) using vegan, gradientForest, lfmm2, tidyverse, and ggplot2, with scripts organized as numbered modules for reproducibility (Aitken et al., 2024).

### M6. Integrated Multi-Omics Analyses

#### M6.1 Stress-Memory Candidate Identification and Prioritization

Stress-memory candidates were identified by integrating seasonal RNA, chromatin accessibility (ATAC-seq), histone modification (H3K4me3 and H3K27ac), DNA methylation (WGBS), and three-dimensional chromatin (Hi-C) data at gene resolution. The analytical rationale follows the established principle that plant stress memory is most robustly associated with persistent or enhanced transcriptional responses following a prior stress episode, together with sustained chromatin features such as H3K4 trimethylation and accessible promoter chromatin (Lämke et al., 2016; Oberkofler et al., 2021; Kappel et al., 2023; Pratx and Bäurle, 2024).

All input layers were converted to a common gene-centred representation across six seasonal time points (m02, m04, m06, m08, m10, m12). Gene-associated RNA, ATAC-seq, H3K4me3, and H3K27ac matrices were derived from deepTools computeMatrix outputs and converted to per-gene seasonal values by averaging bins assigned to each sample block. WGBS data were summarized to gene bodies using the Prosopis genome annotation; for each gene, a length-weighted average methylation value was calculated across overlapping bedGraph intervals, and replicate trees were averaged within each season to produce seasonal CG, CHG, and CHH gene matrices. Hi-C information was either taken from an existing gene-level structural table or derived from seasonal .mcool files as a local cis-contact score in a window centred on the gene midpoint (Abdennur and Mirny, 2020).

Seasonal windows were defined a priori: m02 and m12 as cool-season baseline; m04 as early warming; m06 and m08 as the hot-stress interval; and m10 as the post-stress persistence window. This design targets loci induced during peak heat that retain an altered molecular state after conditions moderate, consistent with the mechanistic definition of stress memory as the maintenance of an altered transcriptional or chromatin state after the inducing stimulus has passed (Oberkofler et al., 2021; Kappel et al., 2023; Pratx and Bäurle, 2024).

For signed layers (RNA, ATAC-seq, H3K4me3, H3K27ac), hot-season induction was quantified as the difference between the hot-season mean and baseline mean; persistence was the retained shift at m10 relative to baseline. For unsigned layers (DNA methylation, Hi-C), change magnitude was used because both gain and loss can represent informative memory-associated states. Per-layer features were combined into a composite memory score, weighting RNA persistence and H3K4me3 persistence most heavily, as these marks are repeatedly implicated in plant heat-stress transcriptional memory (Lämke et al., 2016; Oberkofler et al., 2022; Kappel et al., 2023; Pratx and Bäurle, 2024). DNA methylation was included because stress-associated methylation dynamics can contribute to persistent regulatory states and because histone–methylation crosstalk is documented in plant stress memory (Liu et al., 2014; Oberkofler et al., 2021). Three-dimensional chromatin was included because stress-responsive gene regulation can be conditioned by chromatin topology, and multi-layer integration provides a stronger systems-level framework than single-assay screening (Kappel et al., 2023; Pratx and Bäurle, 2024). Genes passing an initial low-stringency threshold were retained as broad stress-memory-like candidates.

High-confidence candidates were selected by requiring: (i) RNA persistence as mandatory (sustained induction or retained transcriptional competence after peak heat); (ii) at least one corroborating active chromatin layer (persistent ATAC-seq, H3K4me3, or H3K27ac); and (iii) persistence in at least two layers overall. This mirrors the literature criterion that transcriptional memory is distinguished from ordinary stress responsiveness by the combination of persistent transcript levels and sustained chromatin features (Lämke et al., 2016; Oberkofler et al., 2021; Kappel et al., 2023).

InterPro-derived functional annotation was added after filtering by mapping LOC gene identifiers to external protein-to-gene annotation tables. Annotation text was scanned for stress-related keywords spanning heat response, chaperones, ABA signalling, drought, water balance, wax and cuticle biology, reactive oxygen species metaboliM, transport, photosynthetic protection, and chromatin regulation. High-confidence candidates were further intersected with two population-genomic evidence layers: environmental-association SNPs from landscape genomics (prioritizing temperature-related bioclimatic predictors) (Aitken et al., 2024), and XP-CLR outlier genes (Chen et al., 2010). Genes supported by both seasonal molecular memory signatures and population-genomic adaptation evidence were prioritized as plausible links between within-generation stress memory and long-term environmental adaptation. Throughout the manuscript, the full broad set is termed stress-memory-like candidates, and the filtered subset is termed the high-confidence stress-memory candidate set, reflecting the consensus that controlled priming-recovery-retrigger experiments provide the most rigorous demonstration of true stress memory in plants (Oberkofler et al., 2021; Pratx and Bäurle, 2024).

#### M6.2 Regulatory Network Visualization and Cluster Mapping

Season-resolved regulatory networks were constructed from four tab-delimited tables generated in the upstream promoter, motif, and chromatin-loop analysis: network_nodes.tsv, tf_to_distal_edges.tsv, distal_to_promoter_edges.tsv, and tf_to_promoter_edges.tsv. The node table defined unique identifiers, display labels, and gene identifiers for three node types: transcription factors (TFs), distal regulatory anchors, and promoter-associated genes. Three complementary evidence classes were encoded: TF-to-distal links (inferred from motif occurrence in accessible chromatin), distal-to-promoter links (connected via chromatin-loop anchors), and direct TF-to-promoter links, with TF-to-promoter edges emitted only for loops with confirmed promoter-motif support (Grant et al., 2011; Bailey et al., 2015; Rao et al., 2014; Mumbach et al., 2016). Seasons were ordered as m02, m04, m06, m08, m10, and m12.

Interaction tables were aggregated using custom Python scripts: node labels were cleaned by removing technical prefixes (TF::, DISTAL::, GENE::, gene-) to produce manuscript-readable identifiers. TF-to-distal interactions were aggregated by unique TF-distal anchor pair, recording the seasons of support, unique loop identifiers, and maximum and mean edge support scores. Distal-to-promoter interactions were similarly aggregated. Season- and loop-supported TF-distal-promoter triplets were constructed by joining these tables on distal anchor, season, and loop identifier.

Oriented radial network figures were generated with custom Python scripts using pandas and Matplotlib (McKinney, 2010; Hunter, 2007). Because the full regulatory network was too dense for direct plotting, a high-confidence subset was selected based on recurrence, connectivity, and support. Distal anchors were ranked by number of target genes, seasonal breadth, and cumulative edge support; TFs were ranked by promoter and distal targets, seasonal breadth, and total support. In the radial layout, TFs occupied the inner ring, distal regulatory anchors the middle ring, and promoter-associated genes the outer ring. Node sizes were scaled by network degree and node colors by the dominant supporting season; edges were drawn as curved segments colored by season with line width proportional to the square root of the edge_support_score. Final figures were exported as PDF and PNG.

To relate regulatory connectivity to seasonal expression programs, gene lists from heat-positive clusters (Clusters 1 and 2) and the cool-positive cluster (Cluster 3) were mapped onto the full network. Relevant enhancer-promoter edges and direct TF-promoter edges were extracted for each cluster, and TF-distal-promoter triplets were assembled by joining the relevant tables. Results were exported as structured Excel workbooks and used to assess whether heat- and cool-associated expression clusters were preferentially connected to specific TFs, distal regulatory anchors, and seasonally recurrent loops. Statistical and biological interpretation was based on the full interaction and cluster-mapping workbooks rather than on plotted edges, given that the visualization step applies intentional density filtering for readability.

#### M6.3 Post-FIMO Regulatory-Loop Integration and Network Export

This section describes the downstream computational processing performed after seasonal TF-motif-target tables had been generated from FIMO motif-scanning analyses (Grant et al., 2011; Bailey et al., 2015). The workflow integrated loop coordinates, promoter and distal anchors, promoter gene identifiers, motif assignments, candidate TF-family mappings, and quantitative chromatin and RNA measurements from a seasonal expanded TF-motif-target table.

For seasons m04, m06, and m08, which were treated as heat-associated, and m06 and m08 as the hot core, FIMO motif calls were consolidated using a patch script that retained the best site per sequence_name/motif_id combination (lexicographically Mallest q-value then P-value). Expanded seasonal rows were collapsed to one representative record per season and loop_id using collapse_loop_level_summary.py, accumulating promoter gene identifiers, anchor coordinates, motif IDs, motif alternative IDs, motif-family labels, and candidate TF genes as non-redundant comma-separated fields. A loop was flagged as having promoter-motif support when any expanded row linked it to a promoter motif call.

Loop prioritization (rank_high_confidence_loops.py) assigned a total_confidence_score from four components: (i) a binary support score weighting promoter-motif support and evidence for open and active chromatin at promoter and distal anchors; (ii) a motif-richness score capturing distinct motif identifiers; (iii) a candidate-TF score representing inferred TF gene counts; and (iv) a signal-support score from log1p-transformed promoter ATAC, H3K4me3, H3K27ac, RNA coverage, distal ATAC, and distal H3K27ac. Loops were assigned rule-based confidence classes: high_confidence_active (when promoter-motif support, openness, activity, distal openness, distal activity, and promoter RNA signal were all present), moderate_confidence_active, distal_primed, promoter_active_only, and weak_support. Gene-level recurrence (rank_recurrent_loop_gene_candidates.py) aggregated evidence across promoter_gene_ids, tracking seasons, unique loop identifiers, motif diversity, TF support, and the best-supported loop instance. A composite recurrence_score weighted seasonal breadth, hot-core recurrence, and promoter-motif support. Recurrence categories included recurrent_high_confidence, recurrent_moderate_confidence, hot_core_recurrent, broad_season_recurrent, and limited_recurrence. A manuscript-facing ranking script (make_manuscript_ready_heat_candidates.py) computed a heat_priority_score from hot-core season count, heat-season count, total seasons present, TF gene count, motif diversity, and loop confidence, assigning candidates to four priority tiers. Functional annotation was integrated by matching gene identifiers to InterPro and GO terms (Jones et al., 2014; Gene Ontology Consortium, 2026).

Network export scripts (build_loop_network_tables.py) transformed integrated loop summaries into explicit node and edge tables for graph analysis. Reduced subsets were generated by build_reduced_top_edge_network.py based on top-scoring edges or heat-core gene filters. Cytoscape-ready tables with node colors, shapes, and edge styles were generated with export scripts (Shannon et al., 2003). For rapid plotting outside Cytoscape, a NetworkX-based script imported Cytoscape bundle tables, positioned nodes using a force-directed spring layout, and filtered the graph by season, edge type, minimum edge support, or top-edge threshold before drawing (Hagberg et al., 2008; Hunter, 2007). All scripts were implemented in Python 3 and produced plain-text TSV outputs, formatted Excel workbooks, and publication-ready PDF/PNG network figures. Because ranking components and thresholds were explicitly encoded rather than learned statistically, the workflow constitutes a deterministic, rule-based prioritization framework that integrates motif evidence, chromatin state, transcriptional support, and cross-season recurrence.

#### M6.4 Machine-Learning Benchmarking for Seasonal Classification

Seasonal RNA expression profiles spanning six states (m02, m04, m06, m08, m10, m12) were used as the labelled training set. Transcript-level measurements were collapsed to gene level by summing values for identical gene identifiers. Expression values were log2(FPKM + 1)-transformed, and features with no variance across the training set were excluded. Feature panels were assembled from four biological sources: Cluster 1 genes, Cluster 2 genes, Cluster 3 genes (heat-negative), and candidates from stress-memory and bet-hedging analyses. Sample-level summary variables were also generated, including mean cluster scores, eigengene-like first-PC scores, a heat-positive score (Cluster 1 + Cluster 2), a cool-associated score (Cluster 3), and a heat-minus-cool delta.

Two supervised tasks were benchmarked: (i) exact multiclass season classification (six labels) and (ii) binary thermal-state classification (hot: m06/m08; non-hot: remaining seasons). Model families evaluated were: elastic-net multinomial/logistic regression, random forest, linear-kernel SVM, RBF-kernel SVM, shallow multilayer perceptron (sklearn MLP), and a regularized PyTorch multilayer perceptron (TorchMLP) (Zou and Hastie, 2005; Breiman, 2001; Cortes and Vapnik, 1995; Ching et al., 2018).

Robust validation used leave-one-genotype-out outer splits so that all samples from the held-out genotype were excluded from training in each test fold, reducing optimistic inflation from genotype leakage and following recommendations for grouped cross-validation with non-independent data (Roberts et al., 2017). Grouped inner cross-validation was used within each outer training split for model tuning. Performance metrics included accuracy, balanced accuracy, macro F1, weighted F1, Matthews correlation coefficient, Cohen's kappa, log loss, and area under ROC and precision-recall curves (Fawcett, 2006; Saito and RehmMeier, 2015; Chicco and Jurman, 2020). Season-specific diagnostics included top-2 accuracy, adjacent-season accuracy, mean absolute season distance, RMSE of season distance, a weighted season score, heat-likeness RMSE, and a McNemar test comparing the top two exact-season models.

**Table M2.** Benchmarking tasks and principal evaluation metrics.

| **Task** | **Labels** | **Principal Metrics** |
| --- | --- | --- |
| Exact season | m02, m04, m06, m08, m10, m12 | Accuracy, balanced accuracy, macro F1, macro AUROC/AUPRC, top-2 accuracy, mean absolute season distance, heat-likeness RMSE |
| Hot vs. non-hot | Hot = m06/m08; non-hot = m02/m04/m10/m12 | Accuracy, balanced accuracy, macro F1, AUROC, AUPRC, log loss |

### M7. Phenotypic and Biochemical Assays

#### M7.1 Phenotypic and Physiological Measurements

At the end of each simulated seasonal treatment, the following traits were recorded: leaf number, leaflet number, plant height above soil surface, root length, plant diameter, stomatal counts, leaf chlorophyll index, stem moisture status (relative humidity %), soil wetness, soil relative humidity, soil temperature, substrate electrical conductivity (ECp), and soil wet and dry weight. Morphometric traits (plant height, root length, plant diameter, and fresh biomass) were measured manually using a ruler and/or vernier caliper. Stomatal counts were obtained from leaf impressions; images were processed in ImageJ by conversion to 8-bit grayscale, contrast enhancement, and edge detection prior to counting and marking open stomata. Chlorophyll status was measured non-destructively with a SPAD Plus Chlorophyll Meter (Field Scout SPAD 502) and reported as SPAD units. Soil wetness and temperature were measured with a soil moisture meter (Delta T Devices, WET kit and WET-2 sensor), and stem moisture status was recorded using a wood moisture meter (Aneng GN602). Wet and dry soil weights were also recorded for gravimetric soil moisture assesMent.

For the growth-off/metabolic-core-on physiological analysis (Extended Data Fig. 10), observations were grouped by temperature-interval midpoint into lower/mid-temperature and high-temperature classes. For open stomatal fraction, chlorophyll content, and stem relative humidity, bars represent mean interval values with standard error across four intervals per class, while individual interval means are shown as overlaid points. This grouping design follows established frameworks for quantifying stomatal responses under heat stress (Hetherington and Woodward, 2003; Wahid et al., 2007; Santelia and Lawson, 2016).

#### M7.2 Antioxidant Enzyme Assays

Fresh leaf tissue (0.3–0.5 g) was homogenized in 5 ml of ice-cold extraction buffer (100 mM potassium phosphate pH 7.8, 2 mM EDTA, 2% w/v polyvinylpyrrolidone) in a pre-chilled mortar and pestle on ice. The homogenate was centrifuged at 12,000–15,000 × g for 15–20 min at 4 °C, and the supernatant was used as the crude enzyme extract. Where immediate analysis was not possible, aliquots were stored at −80 °C. Soluble protein concentration was determined by the Bradford assay, and all enzyme activities were expressed per milligram of protein.

Catalase (CAT) activity was measured spectrophotometrically at 240 nm. The reaction mixture (4 ml total) contained 2.9 ml of 50 mM phosphate buffer (pH 7.0), 1.0 ml of freshly prepared 10 mM H2O2, and 100 µl of enzyme extract. Absorbance was recorded every 30 s for 180 s, and activity was calculated from the linear decrease in A240 using the extinction coefficient ε = 0.0436 L mmol−1 cm−1 and the formula: Activity (U/ml) = (ΔA240/min × Vtotal) / (ε × d × Vsample), where d is the path length (1 cm).

Peroxidase (POD) activity was assayed using guaiacol as substrate. The reaction mixture (3.0 ml) contained 2.5 ml phosphate buffer (pH 6.1), 0.3 ml guaiacol (20 mM), 0.1 ml H2O2 (10 mM), and 0.1 ml enzyme extract. The increase in A470 due to tetraguaiacol formation was recorded every 30 s for 180 s. Activity was calculated using ε = 26.6 L mmol−1 cm−1: Activity (U/ml) = (ΔA470/min × Vtotal) / (ε × d × Vsample).

Superoxide diMutase (SOD) activity was measured by inhibition of NBT photoreduction. Each 3.0-ml reaction contained 50 mM potassium phosphate buffer (pH 7.8), 9.9 mM L-methionine, 57 µM NBT, 1% Triton X-100, 100 µl enzyme extract, and 1.3 µM riboflavin. Reactions were initiated by riboflavin addition and illuminated for 15 min with two 15-W fluorescent lamps at 8 cm distance. Absorbance was read at 560 nm. Controls lacking enzyme served as maximum-color-development references; non-irradiated tubes served as blanks. SOD activity (U mg−1 protein) was expressed as the amount of enzyme causing 50% inhibition of NBT photoreduction.

For the non-linear antioxidant response analysis (Extended Data Fig. 10), CAT, POD, and SOD activities were modelled across temperature-interval midpoints using degree-2 polynomial regressions with 95% confidence intervals. This non-linear modelling approach was used because antioxidant enzyme activities in plants typically show bell-shaped or threshold-type responses to increasing temperature rather than monotonic stress-collapse trajectories (Wahid et al., 2007; Dong et al., 2018). The polynomial fit and confidence intervals were computed in R using lm() with the formula activity ~ poly(temperature, 2); fitted values and prediction intervals were plotted alongside individual measurements.

#### M7.3 Lignin Quantification

Total lignin content in Prosopis cineraria roots was determined using an acetyl bromide soluble lignin (ABSL) assay. Root tissue was first prepared as alcohol-insoluble residue (AIR) by sequential washing of 100–200 mg of finely ground root powder in 10 ml of 70% ethanol (×3), 10 ml of chloroform:methanol (1:1; ×1-2), and 10 ml of 100% acetone (×1). For each wash, samples were vortexed for 1 min and centrifuged at 4,000 rpm for 5–10 min. Starch was removed by resuspending the pellet in 10 ml of 90% DMSO and incubating for 12–24 h at room temperature (or up to 60 °C for high-starch samples), followed by centrifugation, further ethanol washes, and drying at 40–50 °C in a vacuum oven to constant weight. For lignin solubilization, 5 mg of dried AIR was incubated with 1.0 ml of freshly prepared 25% (v/v) acetyl bromide in glacial acetic acid in a sealed glass vial at 70 °C for 60 min with mixing every 10–15 min. After cooling on ice for 5 min, the sample was diluted with 5.0 ml glacial acetic acid. For spectrophotometric analysis, 0.5 ml of digested supernatant was mixed with 0.5 ml of 2 M NaOH and 0.1 ml of 0.5 M hydroxylamine hydrochloride, brought to a defined final volume with glacial acetic acid, and A280 was measured in quartz cuvettes against a reagent blank. Lignin content was calculated using the Beer-Lambert law: Lignin (g/g AIR) = (A280 × Vtotal) / (ε × l × m), where ε = 23.3 L g−1 cm−1 for woody root tissue, l = 1 cm, and m is the initial AIR mass in grams.

#### M7.4 Epicuticular Wax Quantification

Epicuticular wax content in Prosopis cineraria leaves was determined gravimetrically by rapid solvent extraction. Healthy, fully expanded mature leaves were gently washed with distilled water, air-dried, and 0.5 g of fresh tissue was immersed in 15 ml chloroform for 20–30 s (not exceeding 1 min to minimize extraction of internal lipids). The solvent extract was filtered into a pre-weighed glass vial (W1) and evaporated to dryness at room temperature under a fume hood or at 40–50 °C, then desiccated to remove residual solvent. The vial was reweighed (W2). Epicuticular wax content was expressed on a fresh weight basis as: Wax content (mg/g FW) = (W2 − W1) / fresh tissue weight (g) × 1000.

Seasonal wax content was compared with monthly maximum air temperature using Pearson correlation (Extended Data Fig. 5). Mean wax content per sampling time point was plotted with standard deviation across biological replicates. The correlation between wax content and maximum temperature was calculated from the six paired seasonal means, and the moderately positive relationship was interpreted as evidence of heat-associated wax deposition rather than a strict causal relationship, consistent with published evidence for stress-stimulated wax accumulation in desert-adapted plants (Shepherd and Wynne Griffiths, 2006; Yeats and Rose, 2013). Regulatory support for wax-associated loci was assessed by integrating TF motif and target information with loop-derived promoter-distal interaction annotations as described in M6.2–M6.3.

#### M7.5 Quantitative RT-PCR Validation

Quantitative RT-PCR (qRT-PCR) was used to validate transcript abundance patterns identified by multi-omics analysis and to test candidate regulatory hypotheses across seasonal and experimental contrasts. Target genes included heat stress- and drought-responsive transcripts (heat shock proteins, dehydrins), photosynthesis-associated genes (PEPC, GLK1, RuBisCO), flowering regulators of the FT/HD3A-like family, and the hydrotropiM gene MIZ1. First-strand cDNA was synthesized from 100 ng to 1 µg of total RNA using the iScript cDNA Synthesis Kit (Bio-Rad) in 20-µl reactions (4 µl 5× iScript reaction mix, 1 µl reverse transcriptase, RNA, and nuclease-free water) at 25 °C for 5 min, 46 °C for 20 min, and 95 °C for 1 min. qPCR was performed using SYBR Green PCR Master Mix (Applied Biosystems) in 10-µl reactions containing 5.0 µl 2× master mix, 0.5 µl each primer (10 µM), and 1 µl cDNA template, with an initial denaturation at 95 °C for 10 min followed by 39 cycles of 95 °C for 10 s, 60 °C for 20 s, and 72 °C for 30 s. Melt-curve analysis confirmed amplification specificity. Amplification efficiency for each primer set was verified by five-point serial dilution (90–110% efficiency accepted). Relative transcript abundance was calculated using the comparative Ct (2−ΔΔCt) method, with beta-tubulin as the internal reference gene. Each assay included three biological replicates and three technical replicates, averaged before downstream analysis. Primer sequences and assay details are provided in Table M3.

qRT-PCR data were subsequently integrated with chromatin accessibility, histone modification, and DNA methylation profiles to test regulatory hypotheses, including promoter-proximal enhancer activity and H3K4me3 enrichment at FT and MIZ1, and to evaluate concordance between expression and epigenetic states in progeny as potential indicators of transgenerational stress memory.

#### M7.6 Root HydrotropiM Assays and Lignin Asymmetry Analysis

To investigate the below-ground component of the heat-avoidance response, the expression dynamics and chromatin state of a Prosopis MIZ1-like gene were examined across all six seasonal RNA-seq, H3K4me3 ChIP-seq, H3K27ac ChIP-seq, and ATAC-seq tracks (Extended Data Fig. 6). Concordance of hot-season transcriptional activation with gains in H3K4me3 and chromatin accessibility was evaluated to determine whether the locus carries active chromatin signatures. Seedling root hydrotropiM was assessed under moisture-gradient conditions: seedlings were grown in split-agar chambers in which one agar block was maintained at higher moisture content than the other, and root bending direction toward the moisture source was recorded after 48–72 h. Representative images were annotated for directional bending. Lignin deposition in bent roots was compared between the upper/non-bending region and the lower/bending region using quantitative image analysis of cross-sectional lignin staining or ABSL-equivalent extraction. The two principal anatomical contrasts (PC1-like: upper versus lower, and PC2-like: inner versus outer) were evaluated by pairwise t-test, and P-values are reported for each contrast. Co-expression analysis linked MIZ1 with heat-induced lignin-pathway candidates including 4-coumarate-CoA ligase (4CL), cinnamoyl-CoA reductase (CCR), cinnamyl alcohol dehydrogenase (CAD), laccases, and class III peroxidases, using Pearson correlation of seasonal FPKM profiles. The biological interpretation follows evidence that MIZ1 is required for root hydrotropiM and that laccase- and peroxidase-mediated lignification can asymmetrically alter cell wall mechanics during directional curvature (Kobayashi et al., 2007; Barros et al., 2015; Zhao, 2016).

#### M7.7 Pollen Viability, Phytohormone Profiling, and F2 Validation

Pollen viability was assessed at each seasonal sampling point using fluorescein diacetate (FDA) or acetocarmine staining of freshly collected pollen. The proportion of viable (FDA-positive or strongly stained) grains was scored per slide across at least 200 grains per replicate. Pollen viability was compared across seasonal time points and trees to evaluate whether reproductive performance declined during peak heat and whether genotypes differed in resilience. Variance homogeneity was assessed with Levene's test, and genotype-by-season interaction effects were evaluated using mixed-effects likelihood-ratio models (Extended Data Fig. 7).

Phytohormone abundance was quantified from leaf samples collected at each sampling time point using LC-MS/MS-based methods. Hormones profiled included salicylic acid (SA), abscisic acid (ABA), the ethylene precursor 1-aminocyclopropane-1-carboxylic acid (ACC), and gibberellin-related compounds. Concentrations were log-transformed for visualization and plotted across the seasonal series to characterize SA-to-ABA switching patterns. Independent qRT-PCR validation of heat-cluster genes was performed using the protocols described in M7.5, and the log2 expression response from qRT-PCR was compared with matched RNA-seq values to calculate an overall Pearson concordance across all sampled time points and genes. F2 seedlings derived from the parental tree background were grown under simulated cool-heat cycling in growth chambers (temperature protocols as described in M2). At the end of each simulated seasonal regime, total RNA was extracted and used for RT-qPCR of heat-cluster marker genes. The direction of expression change in F2 seedlings was compared with the direction observed in parental field samples to evaluate recapitulation of the seasonal expression programme. This experimental design avoids over-claiming fixed transgenerational epigenetic inheritance while testing whether stress-responsive transcriptional competence is retained across generations (Philippi and Seger, 1989; Simons, 2011; Crisp et al., 2016; Lämke and Bäurle, 2017).

#### M7.8 Metabolite Quantification and Comparative Leaf Heat-Stress Assay

Malate was quantified from seasonal leaf samples to assess anaplerotic carbon flux associated with the proposed PEPC-MDH-G6PDH metabolic-buffering module (Extended Data Fig. 9). Leaf tissue collected at each sampling time point was immediately flash-frozen in liquid nitrogen and ground to a fine powder. Soluble metabolites were extracted in ice-cold perchloric acid (0.5 M), neutralized with K2CO3, and clarified by centrifugation. Malate concentration in the supernatant was measured enzymatically using malate dehydrogenase (MDH)-coupled spectrophotometric assays or by targeted LC-MS, quantified against a malate standard curve, and expressed as ng/µl per fresh weight equivalent. Samples were grouped into three seasonal classes: cool-transition (February–April), hot (June–August), and cool recovery (October–December). Group means and standard errors were reported, and pairwise Wilcoxon rank-sum tests were used to compare between-group differences given non-normal distributions and Mall group sizes. MDH and glucose-6-phosphate dehydrogenase (G6PDH) FPKM profiles were examined to determine whether expression of these metabolic partners was constitutive or heat-enhanced, in support of the housekeeping anaplerotic model (Chollet et al., 1996; O'Leary et al., 2011; Salvucci and Crafts-Brandner, 2004; Carmo-Silva and Salvucci, 2012).

Leaf thermal stability and chlorophyll retention were evaluated in a detached-leaf heat-stress assay comparing Prosopis cineraria with Medicago sativa as a thermosensitive reference species. Fully expanded leaves were excised and placed in sealed containers at controlled temperatures ranging from 25 °C to 55 °C in 5 °C increments. After heat treatment, leaf chlorophyll was extracted in 80% acetone and quantified spectrophotometrically (A663 and A646 using Lichtenthaler coefficients). Total chlorophyll content was expressed as µg/g fresh weight and plotted against treatment temperature for both species. Lower heat-associated chlorophyll loss in Prosopis relative to Medicago was interpreted as evidence of greater thermostability of the photosynthetic apparatus, supporting the Stay-Green/delayed senescence model consistent with published data on chlorophyll catabolic pathway regulation under heat (Sakuraba et al., 2014).
